# Bioelectronic Modulation of Glioblastoma via Wireless Carbon Nanotube Porin Interfaces

**DOI:** 10.1101/2025.07.04.663176

**Authors:** Fleur E. Groualle, David Onion, Julie A. Watts, Graham A. Rance, Aleksandr Noy, Beth Coyle, Frankie J. Rawson

**Affiliations:** Bioelectronics Laboratory, Division of Regenerative Medicine and Cellular Therapies, School of Pharmacy, Biodiscovery Institute, University of Nottingham, Nottingham NG7 2RD, UK; Flow Cytometry Facility, Medical School, School of Life Sciences, University of Nottingham, NG7 2UH; Nanoscale and Microscale Research Centre (nmRC), Cripps South Building, University of Nottingham, NG7 2RD; Materials Science Division, Lawrence Livermore National Laboratory, Livermore, 94550 USA; School of Natural Sciences, University of California Merced, Merced, CA 94343, USA; Children’s Brain Tumour Research Centre, Translational Medical Sciences, School of Medicine, Biodiscovery Institute, University of Nottingham, Nottingham NG7 2RD, UK

**Keywords:** Membrane Potential, Carbon Nanotube Porins, Electrical Stimulation, Glioblastoma, Cell Cycle, Bioelectricity, Nanomaterials

## Abstract

The membrane potential (V_mem_) and faradaic charge transfer, resulting from altered charge distribution due to ion channels, play a crucial role in cellular bioelectricity. Disruption of V_mem_ can activate pathways associated with cancer proliferation. Manipulating ion channels may therefore present an effective strategy for treating cancers that fail to respond to conventional therapies. One approach to target these channels, is to manipulate the membrane charge which involves the use of wireless bipolar electrodes such as carbon nanotube porins (CNTPs), which could be inserted into cell membranes to mimic these channels. By utilizing membrane dyes, we observed alterations in V_mem_ induced by CNTPs and externally applied electric fields. Analyses of cellular behaviors and processes indicated that V_mem_ is more receptive to stimuli in invasive cancers, while it leads to increased metabolism in less invasive cancers, with notable changes in the cell cycle occurring at approximately 48 hours post-treatment in GBM cell lines. This work shows that CNTPs and electric fields can be used to modulate V_mem_ and alter cancer cell processes, supporting their potential therapeutic capability.

---

All cells exhibit electrical activity, which is responsible for various cellular processes, such as maintaining cellular homeostasis, cell signaling and molecule transport, which play an important role in cancer^1–4^. This electrical activity is also responsible for cellular communication, as cells generate and receive bioelectric signals^5^. This cellular bioelectricity is derived from two sources: (i) ionic currents which are generated by the flow of charged ions, and (ii) faradaic currents arising from the redox reactions of biochemical molecules^2^. Ionic currents can produce endogenous electric fields, as well as promoting diffusion and electrophoresis, as they drive the flux of ions across the plasma membrane^6,7^. Ions diffuse by changing the surface membrane charge distribution through ion channels^8^, which produces the V_mem_^9,10^. Research shows that ion transport influences crucial cellular processes such as excitability, proliferation, the cell cycle, and differentiation^11^, while ion channels help maintain cellular homeostasis by regulating ion flux ^12^. As V_mem_ is essential for ion transport, its modification can affect cell function^11^. The disruption of ion channel expression and flux results in the proliferation, metastasis and metabolic alterations of cancer cells, with various ion transport systems and channels contributing to aerobic glycolysis which is a hallmark of cancer^13–16^.

Glioblastoma Multiformes (GBMs) are grade IV gliomas, highly aggressive brain tumors, with only 6% of patients surviving 5 years or more after their diagnosis^17,18^. Although surgical methods can reduce tumor mass, tumor cells often invade healthy tissue, making complete resection unfeasible^19^. Other methods such as chemotherapy and radiotherapy focusing on the post-surgical residual tumor core have consistently failed to improve survival in clinical trials, as a result, GBMs remain hard-to-treat^18^.

GBMs exhibit a tumor microenvironment that relies on complex cell-cell communication^18^, making them suitable for bioelectric manipulation to regulate cancer signaling pathways and inhibit proliferation ^2^. For instance, glioma cells are characterized by a depolarized resting membrane potential of -20 to -40 mV^20^, and multiple cellular processes such as proliferation, metabolism and migration are attributed to specific ion channels, such as K^+^, Ca^2+^, Na^+^ and Cl^-^ channels ^21^. In support of their importance in GBM, patients with Na^+^ channels mutations experienced a significantly shorter survival time compared to those without those mutations^22^. K^+^, Na^+^ and Cl^-^ channels possess a subset of ion channels known as Voltage-Gated Ion Channels (VGICs,) which are selectively permeable to their respective ions and respond to fluctuations in the membrane potential^23–25^.

Within cancer cells, it has been shown that externally applied electric fields can activate VGICs more readily, enhancing ion flow, which can either make the V_mem_ less negative, known as depolarization, or more negative, known as hyperpolarization, depending on the ion movements and the type of channel(s) activated^26^. Therefore, modulating these channels would provide a safer therapeutic target for tumor regression, as opposed to current aggressive popular treatment such as chemo/radiotherapy, which have many negative side effects which hinders the patient’s quality of life^27–30^. In hard-to-treat cancers, such as GBMs^31^, VGICs significantly influence tumorigenesis^8^, indicating their modulation may offer therapeutic targets for GBMs. One method for this modulation involves electrically communicating with cells^32^.

Previous attempts at electrically communicating with cells were invasive, as they pierced the plasma membrane, disrupting cellular functions and causing perturbations^1,33^. A potential solution is to create biocompatible bioelectronic devices using highly conductive nanomaterials that can be “wirelessly” controlled through bipolar electrochemistry, known as Bipolar Electrodes (BPEs)^1,34^. BPEs can also be fabricated as Biomimetic Proteins (BMPs), which are promising synthetic nanopores that mimic transmembrane proteins, allowing precise molecular transport regulation^3,35^. Additionally, some BMPs can artificially target ion channels for ion selectivity ^36^ reducing ion transport to channels that promote cell growth.

An ideal BMP candidate in this instance is the carbon nanotube (CNT). Short CNTs (<10nm), characterized by a hexagonal lattice of sp^2^-hybridized carbon atoms, can function as a biological analogue protein known as the carbon nanotube porin (CNTP). These CNTPs can mimic biological ion channels by spontaneously integrating into lipid bilayers and cell membranes and have been shown to facilitate rapid water transport. They also exhibit stochastic gating and can form ionic current blockades, without the need for direct physical contact with a power source to regulate ion transport^37^. This intrinsic behaviour enables passive modulation of membrane potential, making CNTPs uniquely suited for bioelectronic applications where external control is impractical or undesirable.

Previous applications of BPEs were polarized in the kV region^38,39^, which is not considered “cell-friendly” and can disrupt cell membranes. In contrast, nano-BPEs function as wireless electrochemical electrodes^40^, with their smaller size allowing targeting of individual cancer cells, they have the capability to modulate bioelectrical effects on cellular behavior, thereby reducing the necessity of surgical intervention^3^.

Recent studies have shown that CNTPs self-insertion into Giant Unilamellar Vesicles (GUVs) can serve as a simplified cellular model, providing a cell-sized confinement to study biochemical reactions and properties of CNTPs ^41,42^. CNTPs have also successfully acted as wireless BPEs, mediating redox processes and influencing cellular behavior, particularly in neuronal cell lines, by polarizing at low biocompatible voltages to modulate electron transfer across membranes^43^, while also being valuable tools in cancer diagnosis and therapy^44^. Building on this, we now explore CNTPs using an externally applied electric field to modulate V_mem_ in GBM cancer cells thereby altering metabolic activity and cell cycle. Our aim is to provide a novel bioelectronic method that, with enhanced development, could branch out to other forms of cancer and be a future, safer treatment in cancer therapies^45^.

In this paper, we explore the application of CNTPs within GBMs, examining how externally applied electric fields, and a combination of the two influence their membrane potential and signal characteristics, as well as their subsequent effects on their metabolism and cell cycle. Figure 1 details the methodology to fulfil these research objects. Figure 1A demonstrates the incorporation of CNTPs into GBM cells via cell culture. Figure 1A details how CNTPs in GBM cells facilitate ion transport. It illustrates how the modulation of ion transport influences cellular bioelectricity, as well as key cellular processes such as metabolic activity and cell cycle progression.

**Figure 1.**
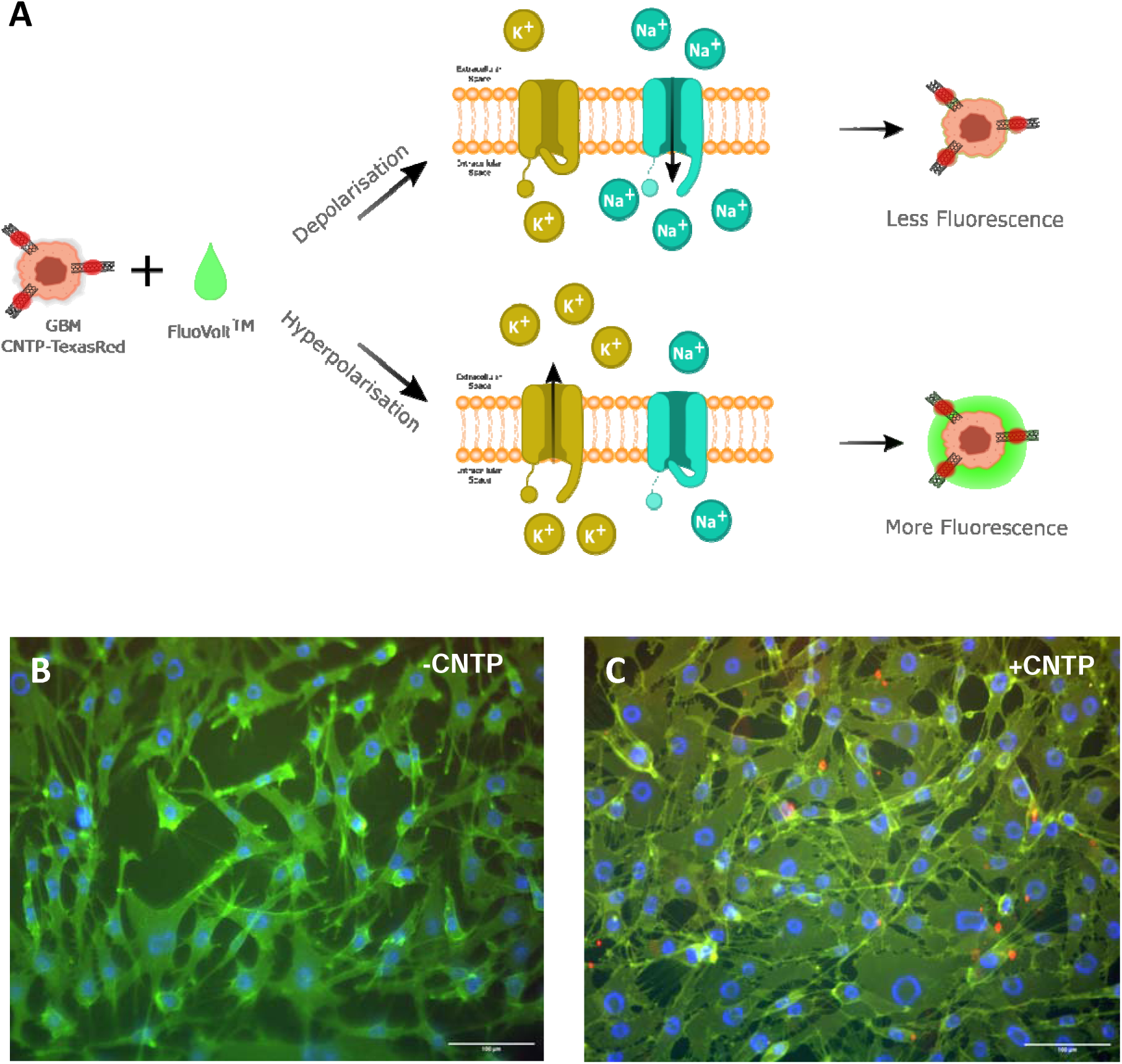
A) Schematic illustrating the mechanism of FluoVolt^TM^ dye fluorescence alteration in response to changes in membrane potential. B) and C) Fluorescence microscopy images of GCE-31 cells with FluoVolt™ in the absence and presence of CNTP-TexasRed, respectively. In B) and C), specific colors correspond to respective Excitation/Emission: FluoVolt™ – 470/525 nm (Green); NucBlue – 350/465nm (Blue); CNTP-TexasRed – 544/581 nm (Red). Scale bar = 100 µm.

## RESULTS & DISCUSSION

We aimed to utilize CNTPs as bipolar electrodes to facilitate charge transfer across the cell membrane. CNTPs were synthesized using previously established methods^46^, then evidence for their inclusion into the cell membrane was established^43^. During the manufacturing process of CNTPs, CNTs are coated with lipid molecules to improve the dispersion of CNT-associated aggregation, increase solubility as well as improving their ability to exhibit /properties of natural membrane-bounded structures, which can be used to increase their compatibility with biological membranes^47–49^.

Evidence of liposomes on the surface of the CNTPs was confirmed using Raman spectroscopy (Figure S1), CNTPs were shown to be present within lipid membranes using cryo-TEM (Figure S2) and a proton translocation assay confirmed that CNTPs affected proton transport ability within lipid membranes (Figure S3).

The GBM cells used in this study were two cell lines isolated from the same patient’s tumor: GCE-31 cells were isolated from the core of the tumor, whereas GIN-31 cells were isolated from the infiltrative margin ^18^. These patient-derived cell lines reflect the heterogeneity of GBM and are highly suitable for studying GBM response to external factors, such as inclusion of CNTPs, applied electric fields (EF), or a combination of the two (CNTP+EF).

The surface of CNTPs was modified by labelling Texas-Red fluorescent dye (CNTP-TexasRed) (Figure S4), which can be seen in GBM cells after a 4-hour incubation period (Figure S5).

Having confirmed the presence of CNTPs within the cells, we then investigated whether membrane potential could be modulated through applied electric fields and/or CNTPs, using the FluoVolt™ membrane potential dye (Figure 1).

The mechanism of action of the FluoVolt™ Membrane Potential probe with the CNTP-TexasRed in GCE-31, as a representation for CNTPs within GBMs, is shown in Figure 1A: as the FluoVolt is a potentiometric dye, it binds to cell membranes and fluoresces according to electrical V_mem_ potential fluctuations in its environment. When the cell membrane is hyperpolarized, greater fluorescence is exhibited, in contrast, any decrease in fluorescence implies the cells are depolarized. The fluorescence values were therefore normalized to reflect changes in intensity. Figure 1B shows that FluoVolt™ (green) can be observed in fluorescence microscopy images after a 4-hour incubation, with Figure 1C providing evidence for both FluoVolt™ and CNTP-TexasRed (red) on the same timescale. The green and red channels were applied to both cells; however, red signals are only visible within the +CNTP image (Figure 1C), with the conjugated CNTP-TexasRed demonstrating that CNTPs have been inserted within the cells. The orange shade (Figure 1C), observed in Figure 1C indicates that CNTPs are present in all cells. However, the large red spots suggest that CNTPs tend to cluster in high concentrations, implying that while they are distributed throughout the cell, they preferentially accumulate in specific regions.

When rat prostate cancer cells undergo electric fields, more specifically a Direct Current (DC), these cells are shown to be galvanotactic and that voltage-gated Na+ channel activity that is upregulated in metastatic cells^50^. Furthermore, the combination of CNTPs and applied DC fields have been shown to modulate neuronal cell behavior through wireless control of membrane electron transfer^43^. We therefore aimed to investigate, whether the external application of a direct current (DC) electric field would also influence the membrane potential and if its combination with CNTPs would influence V_mem_.

Figure 2 demonstrates the membrane potential changes of patient-derived GBM cell lines GCE-31 and GIN-31, with and without the presence of CNTPs across different applied voltages in a raster plot format generated from Figure S6, with yellow-colored spikes showing less fluorescence and the purple color showing greater fluorescence. The smoother gradients indicate a gradual decrease in fluorescence, meaning there are smaller spike events, while the more erratic gradient reflect a more abrupt decrease in fluorescence, indicating larger spike events. All the samples demonstrate incredibly rapid changes in fluorescence within short time frames, with notable spikes attributed to the rapid changes in membrane potential. Studies have shown that cancer cell lines exhibit bioelectrical activity similar to neuronal signals in response to external stimuli, such as CNTPs, the data here shows that GBMs also exhibit similar electrical signalling^51^. A common trend observed here is a decrease in fluorescence over time across all samples, which could mean that all cells, regardless of condition, depolarize with time. However, given that both GCE-31 and GIN-31 cells without any external electric fields and an absence of CNTPS also show a decrease in fluorescence over time, as has been previously reported for other cells^52^, this can likely be attributed to photobleaching.

**Figure 2.**
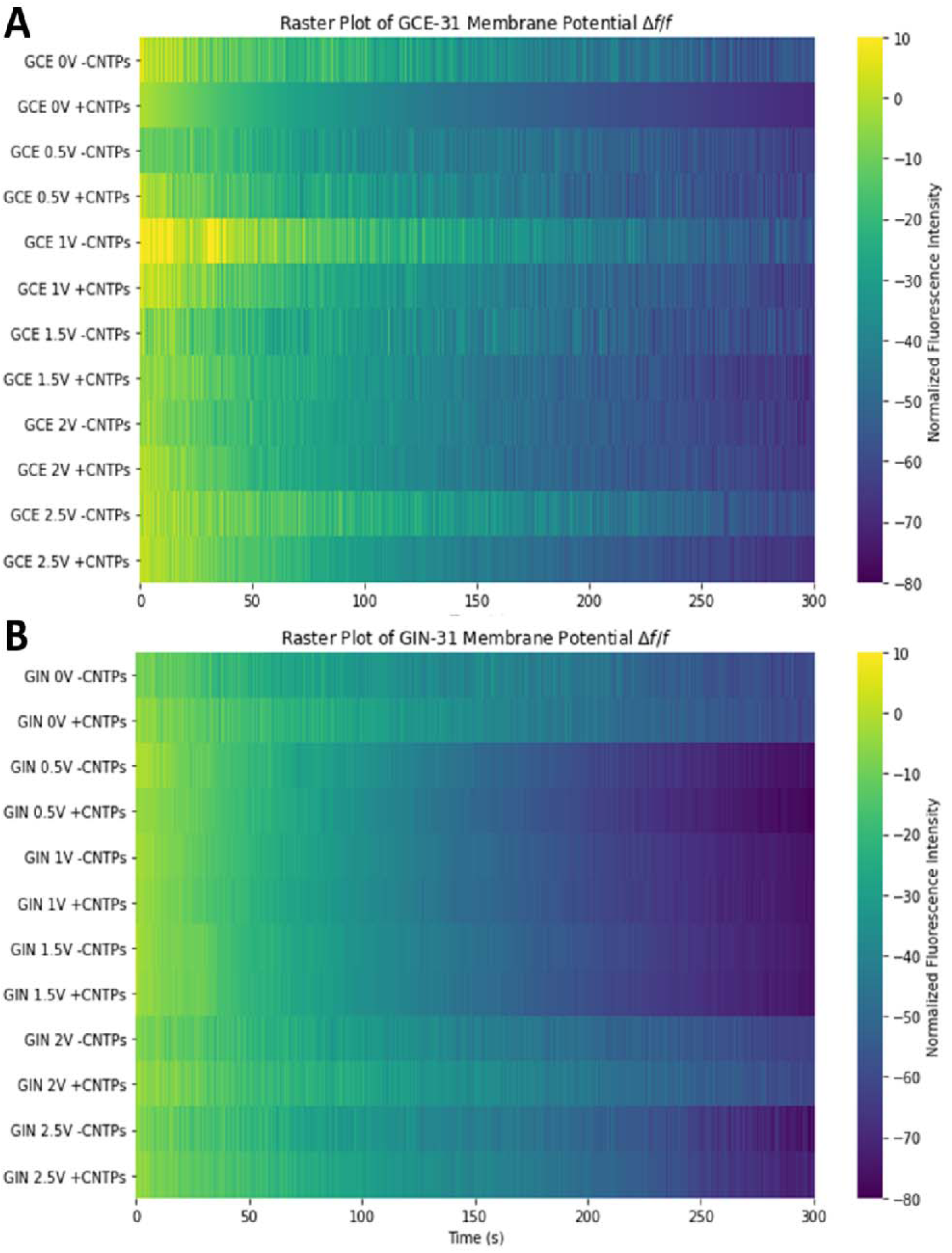
Raster plots of Time-dependent changes in V_mem_ within GBM cells A) GCE-31 and B) GIN-31 with and without CNTP-TexasRed across different applied voltages from 0 to 2.5V spike events denote membrane potential change over time. Normalized cell fluorescence was extracted via ImageJ and used to quantify the percentage change (*ΔF/F*) over time. Biological repeat, N=2, technical repeats, n=10.Colours represent normalized fluorescence intensity, with bright yellow signifying smaller spike changes and darker shades (blue/purple) indicating larger membrane potential spike changes.

### Nonetheless, chacwe can extract valuable data about the behavior of the membrane potential changes

Generally, in Figures 3A and 3B, both GINs and GCEs show that higher voltages (1.5V, 2V, and 2.5V) generate higher-intensity fluorescence signals, with more abrupt color changes, indicating larger spike changes. In contrast, lower voltages (0V and 0.5V) show smaller spike changes, characterized by their smoother gradients. This implies that higher voltages likely trigger more rapid membrane polarizations. GINs appear to have a larger spike magnitude in comparison to GCE-31, however, these are less frequent, appearing “smoother”. Within GCEs, it is evident that in the presence of CNTPs across most voltages, the spikes are smaller in comparison to the analogue in the absence of CNTPs, and the spike magnitude is also more pronounced at higher voltages, suggesting membrane potential change is significantly altered with CNTPs. Within GINs, the opposite is true, as the magnitude is smaller in the presence of CNTPs, therefore, it could be implied that CNTPs stabilize the signal within GIN-31, whereas within GCE-31 this does not occur. Statistical differences of the V_mem_ signal between cells with and without CNTPs at different voltages can be found in Table 1.

**Figure 3.**
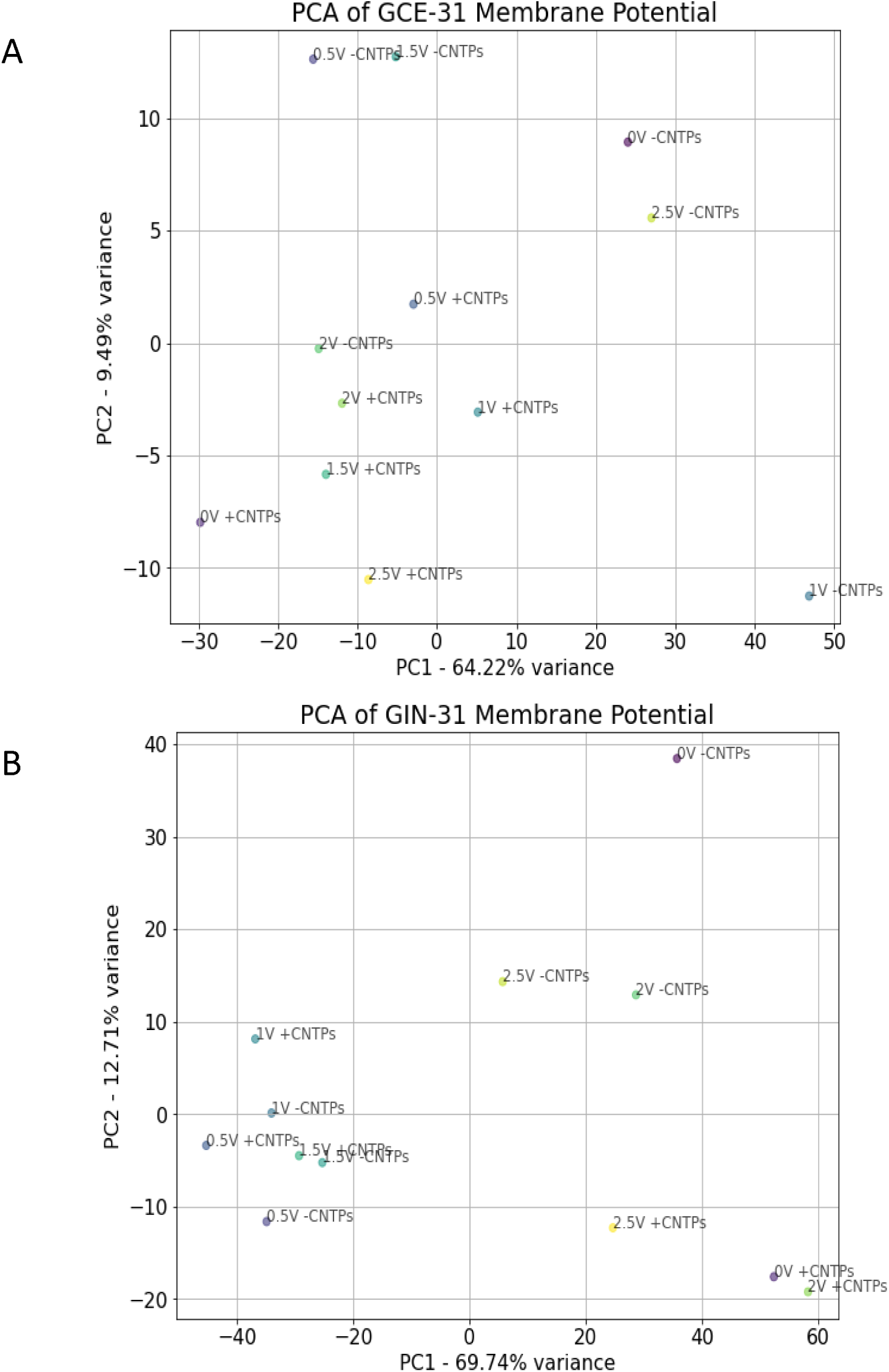
Principal Component Analysis (PCA) of A) GCE-31 and B) GIN-31 generated from Raster plots of Time-dependent changes in V_mem_ (Figure 3). Each point represents a distinct condition of potential, and the absence or presence of CNTPs, which was then standardized with respect to time.

**Table 1.** Statistical results of unpaired t-tests comparing membrane potential of -CNTPs and CNTPs with respect to the applied voltage. ns > 0.05, *p < 0.05, **p < 0.01, ***p < 0.001, ****p < 0.0001.

| Applied Voltage (V) | GCE-31 |  | GIN-31 |  |
| --- | --- | --- | --- | --- |
|  | P-value | Significance | P-value | Significance |
| 0 | 1.44e-27 | **** | 1.24e-06 | **** |
| 0.5 | 2.34e-03 | ** | 0.047 | * |
| 1 | 2.23e-16 | **** | 0.54 | ns |
| 1.5 | 0.079 | ns | 0.44 | ns |
| 2 | 0.53 | ns | 1.23e-12 | **** |
| 2.5 | 2.28e-12 | **** | 8.70e-05 | **** |

GCE-31 and GIN-31 showed significant changes at 0, 0.5 and 2.5V. In the context of 0V, this acts as a control since no electric field is present. This is crucial when comparing membrane potential differences across applied voltages. If no difference is detected between control and CNTP-treated samples at 0⍰V, it becomes difficult to attribute any membrane potential changes observed under applied electric fields solely to the presence of CNTPs. Without a baseline shift, one cannot disentangle whether the CNTPs themselves are mediating the change or whether the electric field alone is responsible. However, in this case, there is a highly significant statistical difference between membrane potential changes in –CNTPs and +CNTPs in both GCE and GIN cells. Therefore, any statistical significance observed in the samples with an applied electric field can be attributed to the fact that even in the absence of an electric field, CNTPs do influence the V_mem_ of GBM cells.

To assist with interpretation, a brief overview of Principal Component Analysis (PCA) is warranted (Figure 3). PCA is an unsupervised dimensionality reduction technique that transforms complex, high-dimensional datasets into a lower-dimensional space while preserving the most significant sources of variance. This approach enables visualisation and quantification of subtle differences that may be obscured in raw data.

Given the high dimensionality of our membrane potential time series data, PCA was employed to extract the key patterns across different treatment conditions. Each data point in Figure 3 represents a single experiment projected in a 2D space defined by the first two principal components (PC1 and PC2), which together capture the majority of the variance in the dataset.

For GCE-31 cells, PC1 and PC2 account for 64.22% and 9.49% of the variance respectively (totaling 73.71%). In this PCA plot, data points with CNTPs cluster separately from those without, indicating that CNTP treatment is a dominant contributor to variance. Increasing applied voltage results in noticeable shifts along PC1 and/or PC2, further separating the conditions. Notably, the 0V condition shows the greatest spread, suggesting high heterogeneity, while 2V samples are tightly clustered, indicating reproducibility under higher field strengths.

In contrast, GIN-31 cells show a higher variance explained by PC1 and PC2 (69.74% and 12.71%, respectively; total 82.45%), implying reduced background noise and greater consistency in membrane potential shifts. Here, conditions such as 0.5V–1.5V with and without CNTPs cluster closely, suggesting less distinction between these treatment groups, while 0V, 2V, and 2.5V conditions show more divergence.

These results demonstrate the utility of PCA in capturing and visualising treatment-dependent patterns in membrane potential responses. The clear separation between CNTP-treated and untreated samples—particularly at higher voltages—supports the conclusion that CNTPs and electric field exposure together modulate membrane potential in a measurable and distinct manner.

Understanding the dynamic behavior of membrane potential changes is crucial for evaluating the effects of external stimuli, such as applied voltage and/or interactions with nanomaterials, on cellular electrophysiology. While absolute membrane potential values provide insight into steady-state conditions, the rate of change = dV/dt, which compares the change in normalized fluorescent intensity, offers a better understanding of how quickly cells respond to external stimuli such as voltage. As CNTPs are known to influence ionic conduction, charge distribution, and electrochemical interactions at the membrane interface, these factors likely contribute to the observed signal effects. To quantify these effects, we calculated the rate of change of the membrane potential signal over time by computing its derivative, visualized in Figure S7. This approach enables us to compare how membrane potential dynamics evolve under different voltage conditions and determine whether CNTPs enhance the responsiveness of the system. Given that bioelectronic interfaces rely on efficient electrical communication between materials and biological systems, assessing the rate of change provides critical insights into how CNTPs modulate electrophysiological behavior, particularly at higher voltages. By employing this analysis, we aimed to determine whether CNTPs accelerate voltage-induced changes in membrane potential, thereby providing a functional measure of their impact on electrical excitability and signal transduction in biological membranes.

Figure 4 demonstrates the peak and trough rates of change for GCEs and GINs. Peaks and troughs were identified, with the rate of change indicating their speed over time extracted from the calculated metrics of the rate of change in Table S2, in addition to the asymmetry index which is used to give insight into the symmetry of the data . It appears that across different voltage conditions, without the addition of CNTPs, that both cell lines respond differently as a function of voltage. The peak activity gradually increased from 0-1V then started falling from 1.5V, eventually reaching below zero at 2V in GCEs (Figure 4A), whereas in GINs the peak activity drops to near or below from 0.5-1.5V before increasing at 2V (Figure 4C). The trough activity is inverse to their peak activity in both cell lines. This implies that there are greater changes observed from 0.5V-1.5V in GCEs, whereas the smallest changes are observed in 0.5-1.5V in GINs, suggesting that this applied voltage range promotes V_mem_ activity in GCEs, whereas it inhibits V_mem_ activity in GINs.

**Figure 4.**
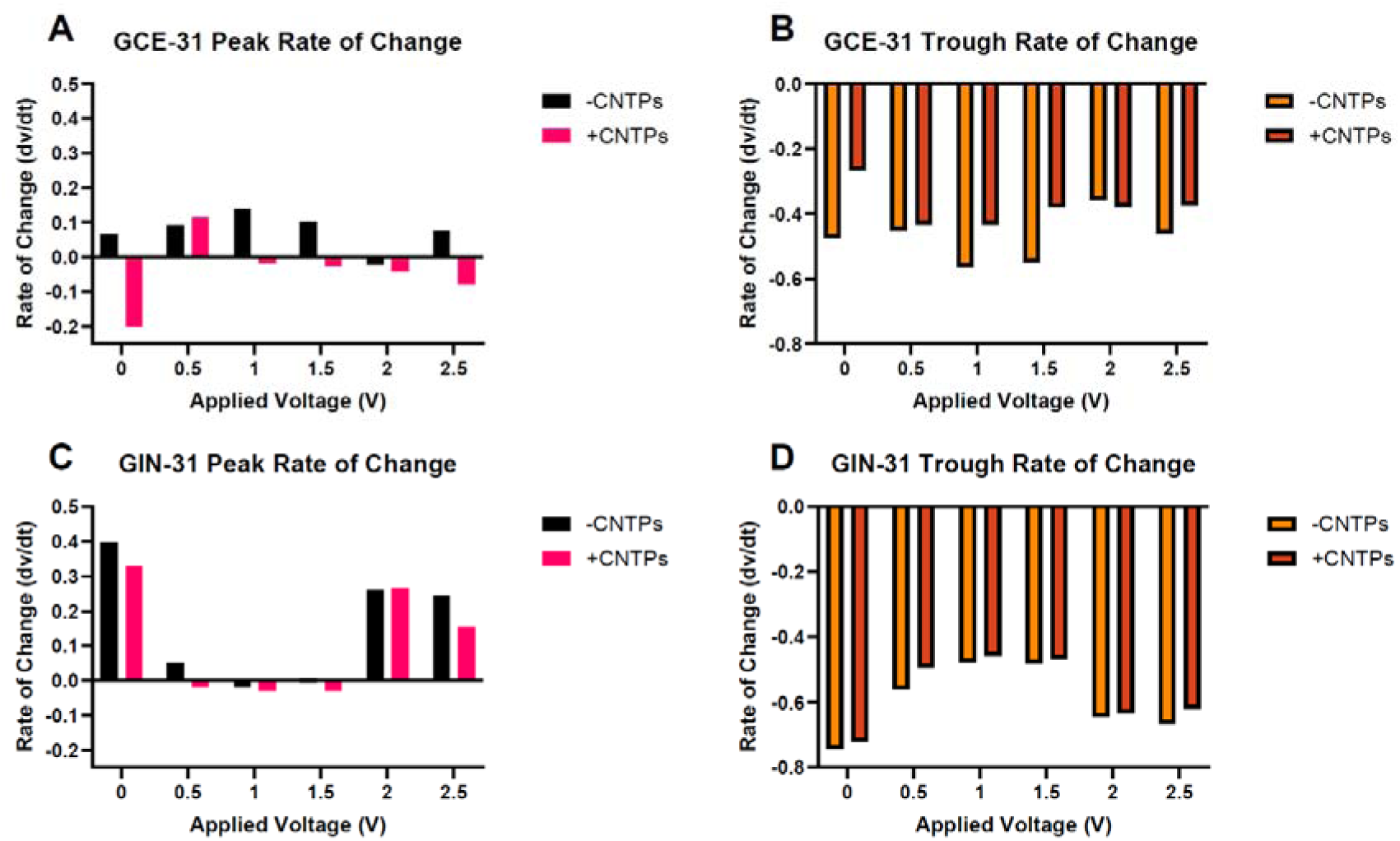
Peak rates of change and trough rates of change metric calculated from Figure 5. A) GCE-31 & B) GIN-31 peak rate of change, respectively. C) GCE-31 and D) GIN-31 trough rates of change, respectively.

Exploring the specific details regarding the extreme values (0V and 2.5V) of the applied voltages may provide insight into the reasons for these observed effects. In Figure 4A, GCE at 0V -CNTP shows a peak rate of change of 0.07, while Figure 4B displays a trough rate of -0.46. To clarify, the asymmetry index quantifies the imbalance between membrane depolarisation and repolarisation rates. An index of 0.14 suggests a moderate imbalance, which can be associated with signal instability and fluctuating membrane potential dynamics. This instability becomes more pronounced at 2.5⍰V without CNTPs, where the peak rate increases to 0.08 and the trough rate modestly improves to -0.46, indicating a steeper upward swing with minimal damping — characteristics of less stable signalling.

In contrast, GIN cells show a more stable profile over the same voltage range. Between 0⍰V and 2.5⍰V –CNTP, the peak rate decreases from 0.39 to 0.24, while the trough rate improves from -0.74 to -0.67, reflecting a reduction in signal volatility and an enhancement in membrane potential stability under increased field strength.

Importantly, CNTPs appear to stabilise the rate of change across both cell lines. In GCE at 0⍰V, CNTPs cause the peak rate to drop from 0.07 to -0.20 and improve the trough rate from -0.46 to - 0.27, reducing oscillations. At 2.5⍰V with CNTPs, the peak rate shifts from 0.08 to -0.08, and the trough rate further stabilises from -0.46 to -0.37, reinforcing this dampening effect.

A similar trend is observed in GIN cells: at 0⍰V, CNTPs reduce the peak rate from 0.40 to 0.33 and improve the trough rate from -0.74 to -0.72. At 2.5⍰V, the peak rate declines from 0.24 to 0.15, and the trough rate stabilises from -0.67 to -0.62.

These findings suggest that CNTPs not only mitigate abrupt shifts in membrane potential but also promote electrical homeostasis, particularly under electrically induced stress. This supports the broader conclusion that CNTPs enhance membrane stability in glioblastoma models by moderating dynamic fluctuations in voltage-dependent activity.

The GIN-31 cells exhibit higher peak rates and lower trough rates than GCE-31 cells, likely due to inherent membrane property differences. At 0V without CNTPs, the peak rate is 0.40, significantly higher than GCE-31, 0.07, while the trough rate of -0.74 indicates stronger oscillations. With CNTPs, the peak rate at 0V reduces to 0.33 and the trough rate to -0.72, stabilizing the signals similar to GCE-31. Increasing the voltage increases the rate of change in membrane potential across both GCE-31 and GIN-31, enhancing ion movement through higher electrical gradients. In GCE-31, the peak rate of change increases from 0.07 at 0V to 0.08 at 2.5V. With CNTPs, these rates shift from -0.20 to - 0.08, emphasizing their voltage-dependent dampening effect. Similarly, in GIN-31, peak and trough rates decrease under different voltage conditions, with CNTPs mitigating fluctuations more effectively than without. Thus, as voltage influences activity, CNTPs modulate these effects in both GCE-31 and GIN-31, likely preventing overstimulation and preserving membrane integrity.

The greatest change in membrane potential correlates with stimulus size - larger stimuli open more ion channels, facilitating greater ion transport and polarization^54,55^.

The observation that CNTPs reduce stimulus amplitude and moderate ion transport dynamics across both GCE-31 and GIN-31 cells suggests a consistent regulatory effect on membrane potential behaviour. Specifically, cells treated with CNTPs exhibit smaller depolarisation events and attenuated repolarisation kinetics, indicating that CNTPs act as modulators of membrane excitability.

This dampening of stimulus response likely reflects a more controlled and stabilised ion flux, possibly due to stochastic gating and ionic current blockade properties inherent to CNTPs, which reduce uncontrolled charge displacement. As a result, CNTPs appear to act as bioelectronic filters, limiting extreme voltage fluctuations while maintaining basal ionic communication.

These findings imply that CNTPs not only alter the rate of membrane potential change but also fine-tune the amplitude of bioelectrical responses, contributing to improved membrane homeostasis. This effect is seen consistently in both GCE-31 and GIN-31 lines, suggesting broad utility of CNTPs across glioblastoma subtypes in modulating bioelectric signalling.

Figure 5 demonstrates that fewer spike events occur in GCEs compared to GINs, indicating an increase in firing activity within GINs. This observation aligns with research indicating that cancers with higher metastatic profiles exhibit greater spiking activity ^51^. GINs are the invasive marginal cancer, suggesting that they possess more significant metastatic properties^19^. Within GCEs, whilst the spiking rate decreases with the presence of CNTPs with no applied voltage, CNTPs across voltages seem to have minimal impact on spiking rates. It could be suggested that GCEs are less sensitive to spiking and that these spike rates may be an intrinsic part of the electrical activity of GCEs compared to GINs, regardless of CNTP presence.

**Figure 5.**
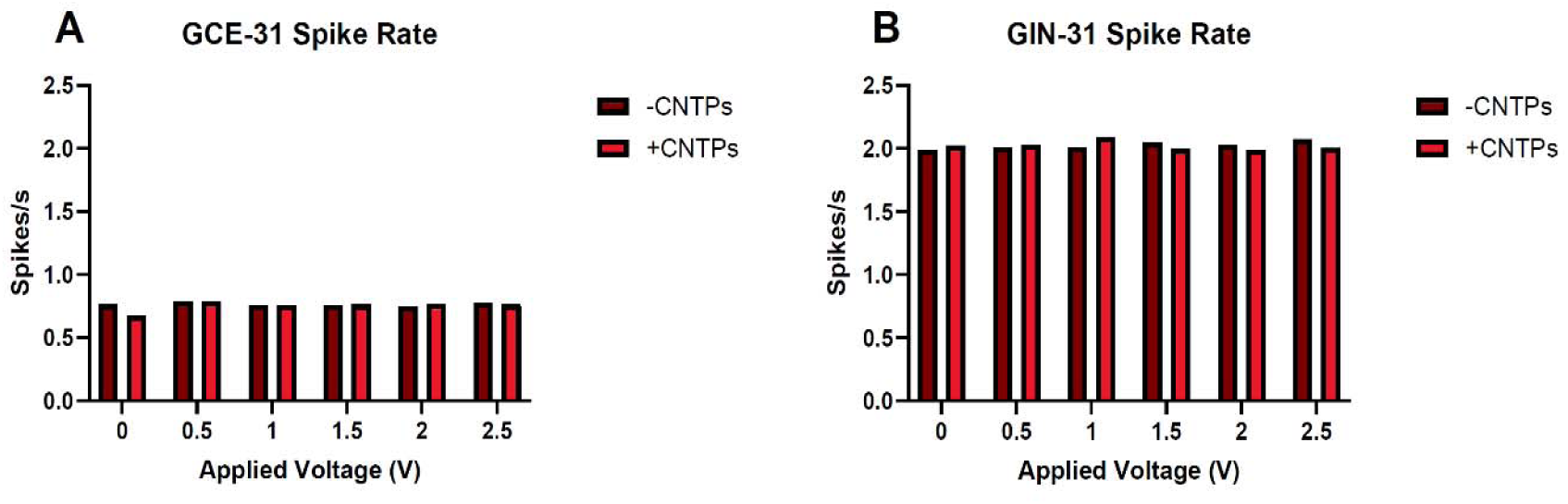
Average spike rates for A) GCEs and B) GINs across applied voltage and absence or presence of CNTPs. The spike rate was calculated using the number of spikes divided by time (seconds), giving a simplified view of the spiking events that are shown in Figure 3.

GINs, on the other hand, exhibit a more notable pattern, as CNTPs amplify spike events at lower applied voltages while inhibiting them at higher applied voltages. This suggests that an absence of electric fields at lower applied voltages with CNTPs facilitates the opening of more ion channels within GINs, while higher applied voltages close them, as research indicates that increased spiking activity is correlated with the opening of a greater number of ion channels^51^. It seems that the spiking events could potentially be modulated by CNTPs, as when a specific threshold is reached (1-1.5V) in GINs since the membrane potential is altered.

To capture the temporal evolution of the frequency component in membrane potential signals, we applied the Short-Time Fourier Transform (STFT), which offers a time-resolved spectral representation of the signal. This analysis differs from the rate of change as it provides a more in-depth examination of the frequency content and its variations over time, in contrast to the rate of change, which focusses on the speed of the signal change over time. It also differs from the raster plots and spike frequency plots found in Figures 3 & 6, respectively, as STFT focuses on the periodic elements of the signal. Figure 6 illustrates that all samples demonstrate an increase in the magnitude of frequency over time. Figures 7bi i-vi shows that the presence of CNTPs generally leads to smoother, more continuous frequency components than in the absence of CNTPs, shown in Figure 6ai i-vi across all applied voltages. This implies that the presence of CNTPs and applied higher voltages within GCEs stabilizes the membrane potential frequency response.

**Figure 6.**
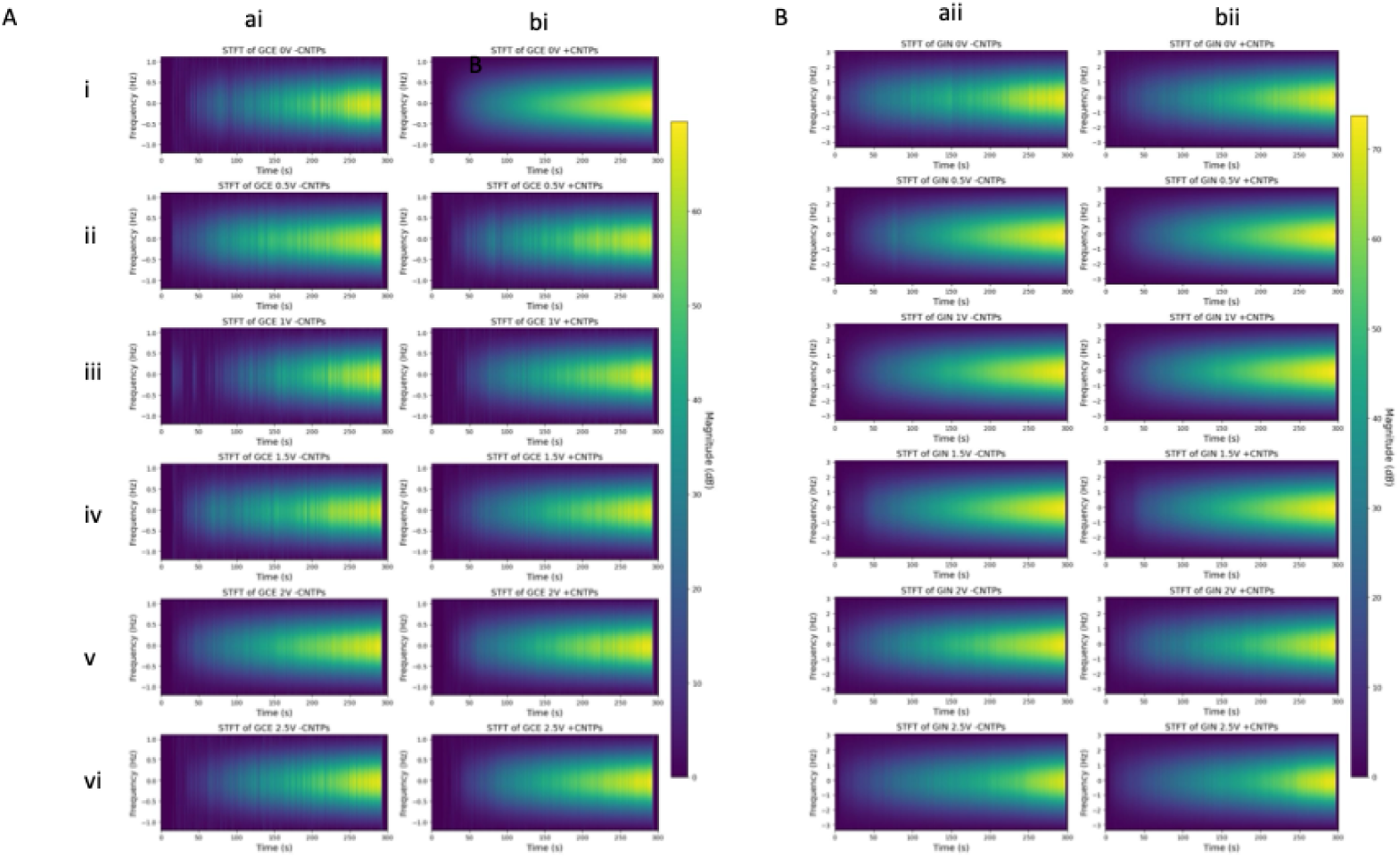
– GCE-31 short-time Fourier transform (STFT) of membrane potential signal at varying applied voltages. GCE cells A-i) 0V, ii) 0.5V, iii) 1V, iv) 1.5V, v) 2V, vi) 2.5V. GCE cells + CNTPs B) i) 0V, ii) 0.5V, iii) 1V, iv) 1.5V, v) 2V, vi) 2.5V. GIN-31 short-time Fourier transform (STFT) of membrane potential signal at varying applied voltages. GIN cells A-i) 0V, ii) 0.5V, iii) 1V, iv) 1.5V, v) 2V, vi) 2.5V. GIN cells + CNTPs B) i) 0V, ii) 0.5V, iii) 1V, iv) 1.5V, v) 2V, vi) 2.5V.

**Figure 7.**
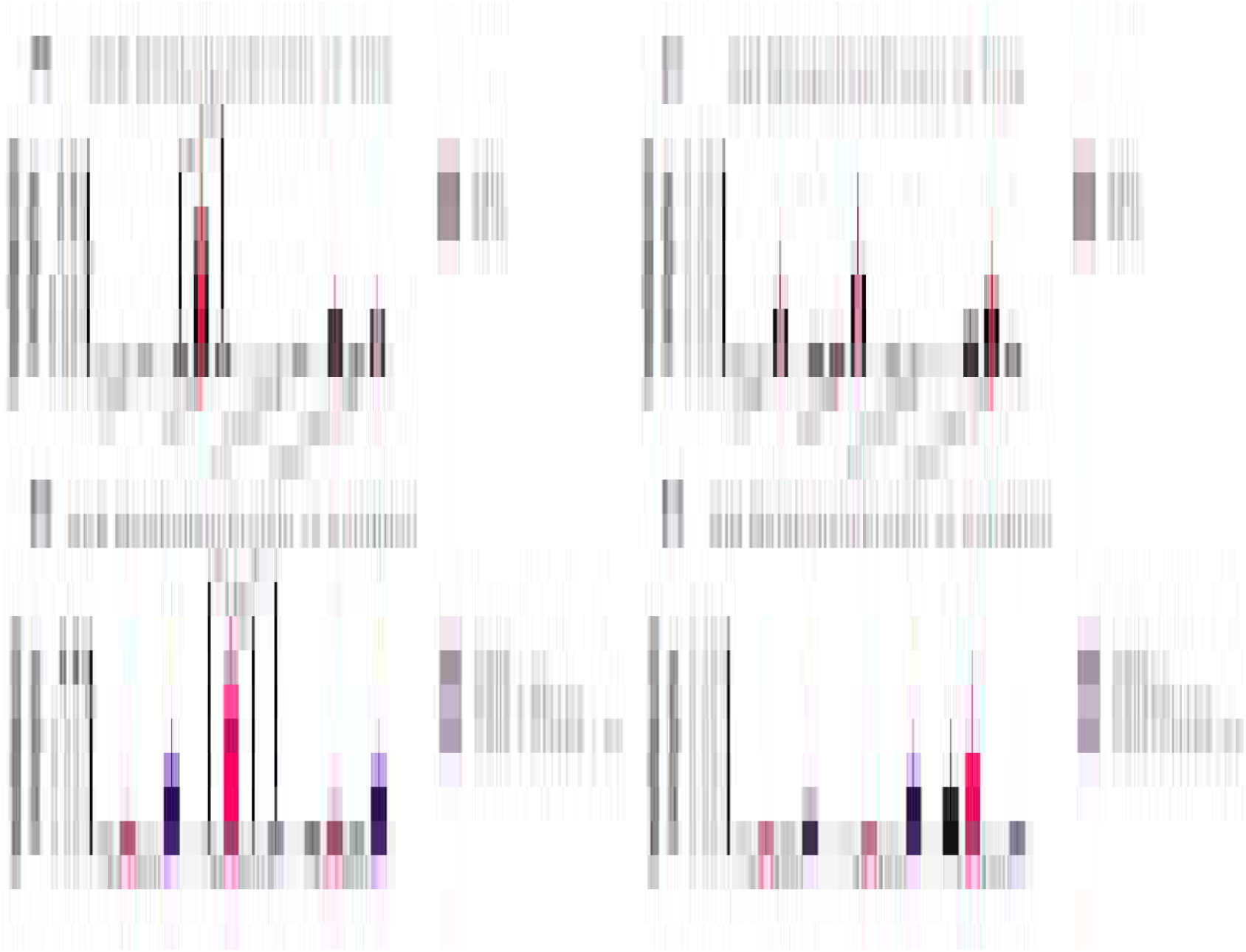
Metabolic activity of GBM Cells, GCE-31 and GIN-31, after 24, 48, and 72 hours under various treatment conditions: untreated, treated with an electric field (EF), treated with CNTPs (CNTPs), and treated with both CNTPs and EF together (CNTPs + EF). ±SEM error bars shown. N=4, n=3.

Figure 6B demonstrates that the STFT analysis of GIN-31 cells shares several similarities with GCE-31 cells as illustrated in Figure 6A, as all samples show an increase in frequency magnitude over time. However, GIN cells exhibit a broader frequency distribution, ranging from -3 Hz to 3 Hz, compared to -1 Hz to 1 Hz in GCE cells. Additionally, GIN cells have a higher magnitude scale (∼70 dB) than GCE cells (∼60 dB) and therefore have a larger frequency range than GCEs. Like GCEs, GINs also show more pronounced ‘yellow’ regions at higher applied voltages, indicating frequency amplification and smoothing in CNTP-containing samples. However, due to their broader frequency range and greater magnitude, GIN cells exhibit a more pronounced response compared to GCE cells. At higher voltages 1.5V–2.5V (Figures 8A&B iv-vi), the yellow regions become increasingly prominent over time, with magnitudes reaching 60 dB or higher, indicating stronger frequency signals in both - CNTP and +CNTP samples. This suggests that increased voltage enhances the magnitude of membrane potential fluctuations in GBM cells, with CNTPs further influencing these dynamics. These findings imply that CNTPs modulate membrane potential differently in GBM cells, with GIN cells demonstrating a broader and stronger frequency response. Other findings have shown that CNTPs possess tuneable ion selectivity as well as transport properties, such as blocking anion transport^37^, suggesting that the observed altered V_mem_ is attributed to this ion selectivity. It is plausible that CNTPs, acting as passive ion-conducting elements, may compete with endogenous ion channels, thereby altering local ion gradients and modulating the amplitude and frequency of potential spikes. This aligns with the observed reduction in membrane potential fluctuations and increased signal stability in both GCE-31 and GIN-31 cells.

**Figure 8.**
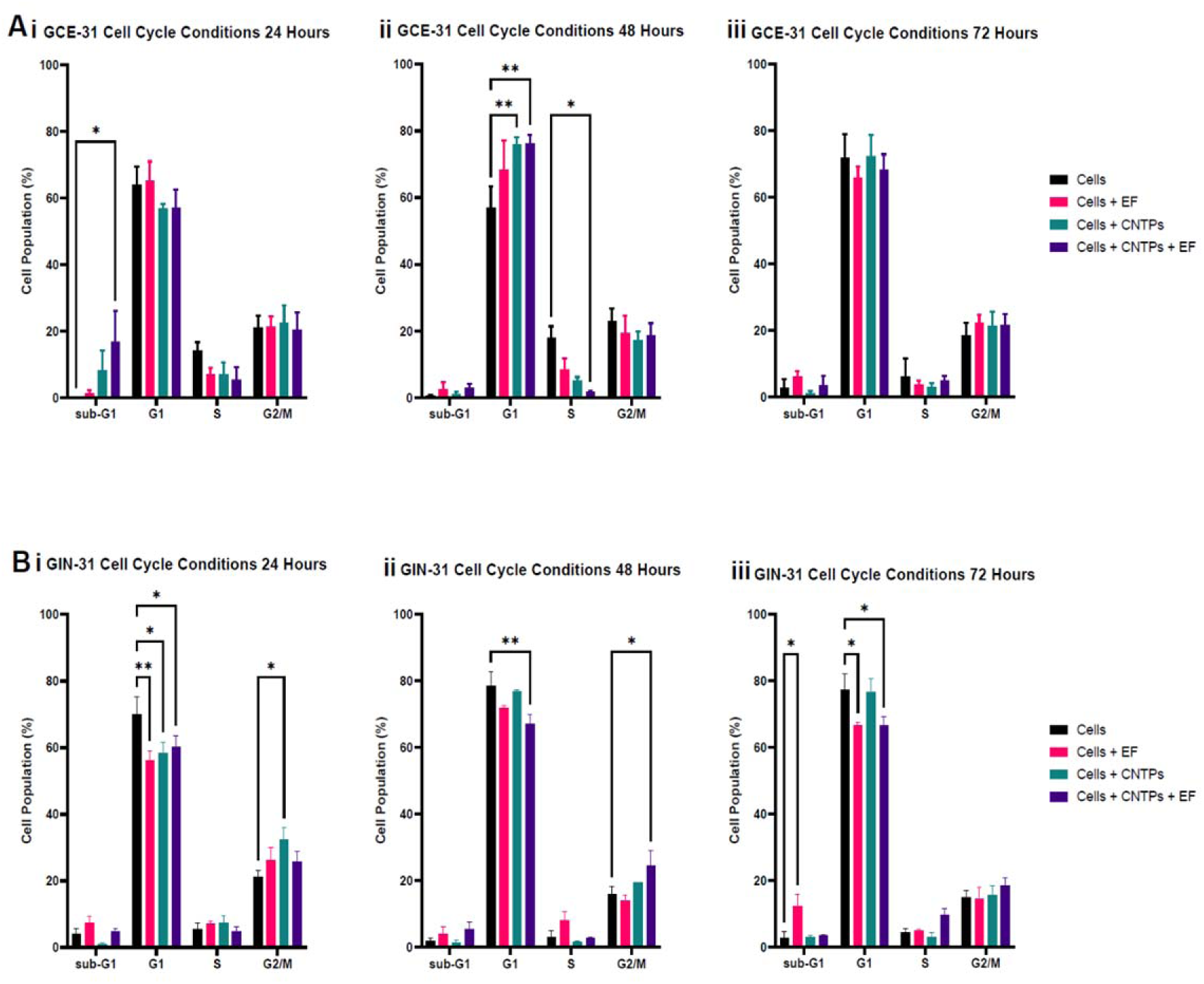
GBM cell cycle response to conditions with/without CNTPs and/or electric fields over time. GCE-31 A i) 24 hours, ii) 48 hours and iii) 72 hours post-treatment, respectively. GIN-31 i) 24 hours, ii) 48 hours and iii) 72 hours post-treatment, respectively. ±SEM error bars shown. N=3, n=3.

Rather than acting as traditional selective ion channels, CNTPs may exert a non-specific, diffusive conductance that partially shunts membrane currents, subtly depolarising the resting potential or flattening dynamic swings. This could explain the attenuated peaks and troughs in voltage change rates and the dampened response to external electric fields.

Thus, while CNTPs may not directly block native ion channels, their integration into the membrane likely disrupts the existing electrochemical landscape, leading to modulated electrophysiological responses. These findings support the interpretation that CNTPs modulate cellular bioelectricity by interfering with native ion flow and stabilising charge distribution, consistent with a bioelectronic modulatory role rather than a purely conductive or channel-like behaviour.

This finding aligns with the PCA analysis found in figure 3, where the GIN-31 dataset exhibited smoother trends, implying greater stability and lower variability.

It is worth mentioning that while the raster plots and STFT plots may appear similar, they provide complementary perspectives. Raster plots focus on discrete spike timing, whereas STFT plots capture spectral details of power distribution across frequencies over time, offering a more comprehensive view of membrane potential stability^35^

To study how the V_mem_ changes caused by CNTPs or CNTPs+EF influence biological processes in GBM cells we first carried out metabolic activity assay (Figure 7). Previous reports have shown that change in V_mem_ is linked to increased/decreased metabolic activity, it has also been reported that cancer cells are susceptible to mitochondria-targeted therapies due to changes in mitochondrial metabolism and mitochondrial membrane potential ^56,57^. Additionally, it has been shown that there is a mechanistic link between cell metabolism, plasma membrane potential and developmental signaling, as changes in cell metabolism alter V_mem_ which then directly influences the activity of the Hedgehog pathway, which controls growth and patterning for insects as well as glycolysis which has an important role in maintaining V_mem_^58^

In this study we found that GCE cells respond to changes in time and conditions with noticeable differences in their metabolic activity (Figures 7A-B), while GIN cells, figure 7C-D consistently have low activity regardless of the experimental setup. For GCE cells, there is a significant increase in metabolic activity after 48 hours when they are exposed to electric fields (EF alone), with absorbance levels higher than in any other condition (p<0.05). This suggests that electric fields enhance metabolic activity in GCE cells. However, adding CNTPs, either by themselves or alongside the electric fields, does not significantly change metabolism at any time point, indicating that CNTPs do not influence metabolic activity in GCE cells in this context. In contrast, GIN cells maintain consistently low metabolic activity, showing no significant difference across all conditions and times, which highlights their lack of response to treatment. Unlike GCE cells, GIN cells do not show any change in metabolic activity over time, remaining inactive whether stimulated by EF or exposed to CNTPs. Overall, GCE cells display higher baseline and metabolic activity compared to GIN cells, pointing to key differences in how they generate energy and respond to EFs and/or CNTPs. These results suggest that while EF can temporarily boost metabolism in GCE cells, CNTPs do not affect metabolic processes in either cell type under these experimental conditions. This indicates that different mechanisms may influence their metabolic activity and how they respond to external factors such as EFs, CNTPs or both.

As aerobic glycolysis is a hallmark of GBM tumours^15^, it could be suggested that GCEs and GINs exhibit different metabolic activity because of how external stimuli due to the V_mem_ differences between GCEs and GINs, as demonstrated in figures 3-7.

48 and 72 under the influence of CNTPs, EFs and CNTPs + EF (**Figure 8**). Within figures 8A&B showed, all conditions showed few sub-G0/G1 events, there was a small increase in sub-G1 events at 24 hours for GCE-31 cells and at 72 hours with GIN-31 cells. At 48 hours GCE-31 cells treated with CNTPS or CNTPs + EF displayed an increase in G0/G1 cells and a simultaneous decrease in those in S phase. In contrast GIN-31 cells showed decreased percentages of cells in G0/G1 in all treatment conditions at 24 hours and simultaneous increase in cells in G2/M cells. When GIN-31 cells are treated with CNTPs + EF phases at 48 hours, there is also a small reduction in cells in G0/G1 phases when treated with EF or CNTPs +EF at 72 hours.

These results indicated that GIN-31 cells are much more responsive to external stimuli than GCE-31, especially with the CNTP + EF conditions and that at 24 hours, the cells are their most sensitive. This is likely related to the doubling time of the cell, which in GIN 31 is ∼24 hours^59^. GCE 31 also demonstrated a doubling time of approximately 24 hours during cell culture.

This cell cycle data can be correlated with the metabolic activity in figure 8. The sharp drop in metabolic activity in GIN-31 cells treated with CNTP+EF aligns with the rise in the sub- G0/G1 population at the same times. This implies the reduced metabolic activity may be caused by increased apoptosis either alone or in combination with other disruption to cell health or cell cycle.

In GIN-31 cells, the effects of CNTPs alone or EF alone are less clear, showing moderate levels of metabolic activity. However, when CNTP and EF are combined, these effects are intensified, suggesting that the treatments work in parallel to speed up metabolic disruption. Interestingly, the GCE-31 cells maintain stable metabolic activity across conditions and don’t show significant changes in their sub-G1 populations. This stability indicates that GCE-31 cells are better able to withstand metabolic and apoptotic challenges.

The differences in responses between GCE-31 and GIN-31 show that GIN-31 cells are more sensitive to energy and death signals from treatments. In GIN-31 cells, lower metabolic rates may hinder their ability to recover or fix damage caused by EF, leading to a larger proportion of apoptotic cells as indicated by the increase in sub- G0/G1 cells. On the other hand, GCE-31 cells keep their metabolic balance, demonstrating that GCEs have inherently different ion channel expressions which are affected by CNTPs, EF and CNTP+EF.

By looking at the data on metabolic activity along with the sub- G0/G1 rates, it becomes clear that decreased metabolism is an early warning sign and a contributor to cell death in sensitive cell lines like GCE-31, especially when facing the combined effects of CNTP and EF treatments at 48 hours. This relationship highlights how CNTPs, EF and CNTP+EF can affect energy metabolism and potentially promote cell death at the same time, with significant effects depending on the cell line.

It is apparent that CNTPs, when combined with an electric field, exert a more significant influence on GIN-31 cells compared to GCE-31 cells, as illustrated in Figures 5, 6, 7, 9 and 10. The enhanced stabilizing effect of CNTPs in GIN cells across voltages, likely relates to their impact on membrane potential and the subsequent effects on the cell cycle, potentially leading to apoptosis.

During the cell cycle, GIN-31 in particular, exhibits significant differences with an increase in the sub- G0/G1 and G0/G1 phases when electric fields and CNTPs are applied. Cancer cells depend on hyperpolarization during the G0/G1 and S phase transition, while they rely on depolarization for the S/G2/M transition^56^. Blocking voltage-gated hERG channels has been demonstrated to result in cell cycle arrest in the G0/G1 phase^31^. Similarly, the blockage of Ca^2+^-activated K^+^ channels (BK) has causes cells to be arrested in the G0/G1 phase^21,31,56,60^. Therefore, it can be inferred from the literature that hyperpolarization observed here may be related to the inhibition of hERG and/or BK K^+^ channels. It could be argued that CNTPs may be considered as passive K^+^ and proton channels in the membrane. Furthermore, the disruption of Cl^-^ channel in the G_2_/M phase prevents any further ion flux^21^. Hence, the presence of electric fields and CNTPs in conjunction with these fields appears to influence the cell cycle. These findings suggest that CNTPs modulate membrane potential dynamics not by mimicking ion channels, but rather by introducing non-specific ionic conductance that competes with, or shunts, endogenous ion flux, thereby dampening sharp transitions in potential and smoothing electrophysiological responses. This passive modulatory role explains the increased stability observed in both cell lines and supports the notion of CNTPs as bioelectronic regulators of cellular excitability.

In support of the hypothesis that CNTPs are impacting the voltage, preliminary V_mem_ data suggests that at 2V, there is a statistically significant difference between GIN samples with and without CNTPs, although this distinction is not observed in GCEs. Metastatic progression is the most lethal aspect of cancer and requires cells to find ways to mechanically adapt ^61^. Since GINs were isolated from the infiltrative margin they are likely to have undergone mechanical adaptations, understanding these may provide insight into why GINs are more receptive to CNTP and applied EFs compared to GCEs.

Overall, it appears that externally applied voltages and CNTPs, do have an impact on V_mem_ within GINs and GCEs, however, it seems that the modulation that occurs is significantly more apparent within GINs. CNTPs and electric fields do enhance metabolism at the 48-hour mark, but this effect is observed exclusively in GCEs. The cell cycle is influenced by CNTPS and electric fields, particularly in GINs.

The presence of CNTPs and an electric field, along with their combination resulting in an induced K^+^ and/or proton flux, and voltage-dependent insertion of CNTPs in the membrane, play a role in influencing the membrane potential, which impacts the metabolism and cellular functions of GCE and GIN cell lines.

## METHODS

### Cell Culture

GBM cells−Glioma INvasive Marginal 31(GIN) cells from the infiltrative tumor margin and GliomaCore Enhanced 31 (GCE) from the core of the tumor were isolated previously from a patient who underwent surgery at the Queen’s Medical Centre, University of Nottingham (Nottingham, UK)^19^. GIN and GCE cells were cultured in DMEM (Gibco) supplemented with 10% FBS, 1% penicillin/streptomycin, and 1% L-glutamine. Cells were maintained at 37 °C in a humidified incubator containing 5% CO_2_. Cells were tested for mycoplasma every month, where they were grown in an antibiotic-free medium for 1 week before mycoplasma testing. All cells used were mycoplasma-free.

### Electrical Stimulation

Cells were seeded at 1×10^5^ cells/well in a 24 well plate (Costar), left to incubate overnight. The following day, the media was aspirated from the well to be replaced 20µL of CNTP-Texas Red or CNTPs and 80µL of DMEM then incubated for t of minimum 4 hours. A direct current (DC) field was applied to cells through two needle feeder electrodes (25G, BD Microlance) in a sanitized modified well plate lid with a distance of 1cm from a Programmable DC Power Source EL-R Series Power Supply (Aim-TTi) and stimulated from voltages 0.5V-2.5V at varying time periods.

### Membrane Potential Imaging

GBM Cells were seeded in 24 well plates at a density of 1 x 10^5^ cells/well incubated at 37°C for 24h. 20µL of CNTP-TexasRed was then incubated in selected cells for 4h. After washing with PBS, the FluoVolt™ Membrane Potential Kit (F10488, ThermoFisher Scientific) was used according to an adapted industrial protocol of using half the reagents for half the incubation period, along with 1 drop of Hoechst 33342 (NucBlue Live ReadyProbes Reagent, R37605, Thermo Fisher Scientific) per well. The CNTPs were aspirated and washed with PBS twice, finally adding 300µL of the FluoVolt and

NucBlue mix to the well and left at room temperature shielded from light. After 10 minutes, the dyes were removed, cells were washed with PBS and 300µL of phenol-free media added. Microscopy images were acquired with a Nikon fluorescent microscope (Model: TI-DH) with an optiMOS sCMOS camera and utilizing NIS-Elements (version 4.60) software. Image sequences of the Stimulation protocol with FluoVolt were captured using a GFP channel, 2×2 binning every 10ms for 5 minutes.

Membrane charging potential was analyzed using Corrected Total Cell Florescence (CTCF) via ImageJ, by analyzing 10 cells within the image, which was then normalized for percentage change (ΔF/F) over time. Additional analyses were performed using Python (Spyder 4.1.5).

### Protein Concentration Assay

Following Stimulation, wells were washed three times with PBS by gentle swirling for 5 min. 500µL of 2% Triton X-100 solution in PBS buffer was then added to each well and the plates incubated 20 min. The lysed cell samples were then transferred to a micro-centrifuge tube and centrifuged at 16000 ×g for 20 min. The supernatant was removed and transferred to a fresh micro-centrifuge tube before analysis using a Pierce™ BCA Protein Assay Kit (A55864, ThermoFisher Scientific) using the manufacture’s recommended protocol. Protein quantification was derived from absorbance values at 562 nm, which was then converted to protein quantity using a standard curve of diluted bovine serum albumin (BSA) standard, ranging from 0-2000 μg/mL. The standard curve was performed N=8 times and averaged to ensure its consistency with all samples for improved accuracy.

### Metabolic Activity Assay

Following Stimulation, WST-1 Assay Reagent (ab155902, abcam) was applied to cells according to manufacturer’s recommended protocol. The Normalized Metabolic activity was calculated from the metabolism abs value/ protein quantification number of the sample.

### Cell Cycle Analysis using Flow Cytometry

Post stimulation, cells were washed with PBS, following which 250 µL of Trypsin was added, and samples were centrifuged at 300 x g for 5 min. After supernatant removal and PBS wash, cells were fixed with 70% ethanol. For flow cytometry analysis the samples were centrifuged at 300 x g for 5 min. The ethanol was then aspirated, and the cells were washed again with PBS. The cells were resuspended in 500 µL FxCycle™ PI/RNase Staining Solution (F10797, ThermoFisher) and subsequently transferred to flow cytometry (FCM) tubes (Greiner, 115101). Cell cycle fluorescence was measured with an ID7000 Spectral Flow Cytometer. Kaluza software (v.2.1) was used to analyze cell cycle data.

## Author Contributions

FEG: Conceptualization, Methodology, Validation, Investigation, Visualization, Formal analysis, Writing—original draft/ review & editing

DO: Flow Cytometry Training and Analysis, Writing - review & editing

JAW: TEM Image Acquisition, Writing - review & editing

GR: Raman Spectroscopy Data Analysis and Acquisition, Writing - review & editing

BC: Supervision, Writing - review & editing

FJR: Conceptualization, Methodology, Validation, Writing – original draft/ review & editing, Supervision, Funding Acquisition.

## Acknowledgements

The authors would like to acknowledge Professor Aleksandr Noy from the Lawrence Livermore National Laboratory for his assistance and support. We would also like to thank Akhil Jain for his help in the microscopy work and reviewing drafts. We also thank Akhil Jian for support in microscopy work and training.

## Notes

The authors declare no competing financial interest.

## REFERENCES

(1) Sanjuan-Alberte, P.; Jain, A.; Shaw, A. J.; Abayzeed, S. A.; Domínguez, R. F.; Alea-Reyes, M. E.; Clark, M.; Alexander, M. R.; Hague, R. J. M.; Pérez-García, L.; Rawson, F. J. Wireless Nanobioelectronics for Electrical Intracellular Sensing. ACS Appl Nano Mater 2019, 2 (10). 10.1021/acsanm.9b01374.

(2) Robinson, A. J.; Jain, A.; Sherman, H. G.; Hague, R. J. M.; Rahman, R.; Sanjuan-Alberte, P.; Rawson, F. J. Toward Hijacking Bioelectricity in Cancer to Develop New Bioelectronic Medicine. Advanced Therapeutics. 2021. 10.1002/adtp.202000248.

(3) Gibney, S.; Hicks, J. M.; Robinson, A.; Jain, A.; Sanjuan-Alberte, P.; Rawson, F. J. Toward Nanobioelectronic Medicine: Unlocking New Applications Using Nanotechnology. Wiley Interdisciplinary Reviews: Nanomedicine and Nanobiotechnology. 2021. 10.1002/wnan.1693.

(4) Sanjuan-Alberte, P.; Alexander, M. R.; Hague, R. J. M.; Rawson, F. J. Electrochemically Stimulating Developments in Bioelectronic Medicine. Bioelectron Med 2018, 4 (1), 1. 10.1186/S42234-018-0001-Z.

(5) Brook Chernet, M. L. Endogenous Voltage Potentials and the Microenvironment: Bioelectric Signals That Reveal, Induce and Normalize Cancer. J Clin Exp Oncol 2014, s1 (01). 10.4172/2324-9110.s1-002.

(6) Funk, R. H. W. Endogenous Electric Fields as Guiding Cue for Cell Migration. Frontiers in Physiology. 2015. 10.3389/fphys.2015.00143.

(7) Payne, S. L.; Levin, M.; Oudin, M. J. Bioelectric Control of Metastasis in Solid Tumors. Bioelectricity 2019, 1 (3). 10.1089/bioe.2019.0013.

(8) Rao, V. R.; Perez-Neut, M.; Kaja, S.; Gentile, S. Voltage-Gated Ion Channels in Cancer Cell Proliferation. Cancers. 2015. 10.3390/cancers7020813.

(9) George, L. F.; Bates, E. A. Mechanisms Underlying Influence of Bioelectricity in Development. Frontiers in Cell and Developmental Biology. 2022. 10.3389/fcell.2022.772230.

(10) Levin, M. Molecular Bioelectricity: How Endogenous Voltage Potentials Control Cell Behavior and Instruct Pattern Regulation in Vivo. Mol Biol Cell 2014, 25 (24). 10.1091/mbc.E13-12-0708.

(11) Sundelacruz, S.; Levin, M.; Kaplan, D. L. Role of Membrane Potential in the Regulation of Cell Proliferation and Differentiation. Stem Cell Rev Rep 2009, 5 (3). 10.1007/s12015-009-9080-2.

(12) Serrano-Novillo, C.; Capera, J.; Colomer-Molera, M.; Condom, E.; Ferreres, J. C.; Felipe, A. Implication of Voltage-Gated Potassium Channels in Neoplastic Cell Proliferation. Cancers. 2019. 10.3390/cancers11030287.

(13) Horvat, A.; Muhič, M.; Smolič, T.; Begić, E.; Zorec, R.; Kreft, M.; Vardjan, N. Ca2+ as the Prime Trigger of Aerobic Glycolysis in Astrocytes. Cell Calcium 2021, 95. 10.1016/j.ceca.2021.102368.

(14) Meyer, D. J.; Díaz-García, C. M.; Nathwani, N.; Rahman, M.; Yellen, G. The Na+/K+ Pump Dominates Control of Glycolysis in Hippocampal Dentate Granule Cells. Elife 2022, 11. 10.7554/eLife.81645.

(15) Alfardus, H.; Mcintyre, A.; Smith, S. MicroRNA Regulation of Glycolytic Metabolism in Glioblastoma. BioMed Research International. 2017. 10.1155/2017/9157370.

(16) Leng, S.; Zhang, X.; Wang, S.; Qin, J.; Liu, Q.; Liu, A.; Sheng, Z.; Feng, Q.; Hu, X.; Peng, J. Ion Channel Piezo1 Activation Promotes Aerobic Glycolysis in Macrophages. Front Immunol 2022, 13. 10.3389/fimmu.2022.976482.

(17) Salari, N.; Fatahian, R.; Kazeminia, M.; Hosseinian-Far, A.; Shohaimi, S.; Mohammadi, M. Patients’ Survival with Astrocytoma After Treatment: A Systematic Review and Meta-Analysis of Clinical Trial Studies. Indian Journal of Surgical Oncology. 2022. 10.1007/s13193-022-01533-7.

(18) Smith, S. J.; Diksin, M.; Chhaya, S.; Sairam, S.; Estevez-Cebrero, M. A.; Rahman, R. The Invasive Region of Glioblastoma Defined by 5ALA Guided Surgery Has an Altered Cancer Stem Cell Marker Profile Compared to Central Tumour. Int J Mol Sci 2017, 18 (11). 10.3390/ijms18112452.

(19) Smith, S. J.; Rowlinson, J.; Estevez-Cebrero, M.; Onion, D.; Ritchie, A.; Clarke, P.; Wood, K.; Diksin, M.; Lourdusamy, A.; Grundy, R. G.; Rahman, R. Metabolism-Based Isolation of Invasive Glioblastoma Cells with Specific Gene Signatures and Tumorigenic Potential. Neurooncol Adv 2020, 2 (1). 10.1093/noajnl/vdaa087.

(20) Bordey, A.; Sontheimer, H. Electrophysiological Properties of Human Astrocytic Tumor Cells in Situ: Enigma of Spiking Glial Cells. J Neurophysiol 1998, 79 (5). 10.1152/jn.1998.79.5.2782.

(21) Elias, A. F.; Lin, B. C.; Piggott, B. J. Ion Channels in Gliomas—From Molecular Basis to Treatment. International Journal of Molecular Sciences. 2023. 10.3390/ijms24032530.

(22) Joshi, A. D.; Parsons, D. W.; Velculescu, V. E.; Riggins, G. J. Sodium Ion Channel Mutations in Glioblastoma Patients Correlate with Shorter Survival. Mol Cancer 2011, 10. 10.1186/1476-4598-10-17.

(23) Bezanilla, F. Voltage-Gated Ion Channels; 2007; pp 81–118. 10.1007/0-387-68919-2_3.

(24) Catterall, W. A. Structure and Function of Voltage-Gated Ion Channels. Annual Review of Biochemistry. 1995. 10.1146/annurev.bi.64.070195.002425.

(25) Armstrong, C. M.; Hille, B. Voltage-Gated Ion Channels and Electrical Excitability. Neuron. 1998. 10.1016/S0896-6273(00)80981-2.

(26) Yang, M.; Brackenbury, W. J. Membrane Potential and Cancer Progression. Frontiers in Physiology. 2013. 10.3389/fphys.2013.00185.

(27) Lang, F.; Stournaras, C. Ion Channels in Cancer: Future Perspectives and Clinical Potential. Philosophical Transactions of the Royal Society B: Biological Sciences. 2014. 10.1098/rstb.2013.0108.

(28) Bortner, C. D.; Cidlowski, J. A. Ion Channels and Apoptosis in Cancer. Philosophical Transactions of the Royal Society B: Biological Sciences. 2014. 10.1098/rstb.2013.0104.

(29) Altamura, C.; Greco, M. R.; Carratù, M. R.; Cardone, R. A.; Desaphy, J. F. Emerging Roles for Ion Channels in Ovarian Cancer: Pathomechanisms and Pharmacological Treatment. Cancers. 2021. 10.3390/cancers13040668.

(30) Sherman, H. G.; Jovanovic, C.; Abuawad, A.; Kim, D. H.; Collins, H.; Dixon, J. E.; Cavanagh, R.; Markus, R.; Stolnik, S.; Rawson, F. J. Mechanistic Insight into Heterogeneity of Trans-Plasma Membrane Electron Transport in Cancer Cell Types. Biochim Biophys Acta Bioenerg 2019, 1860 (8). 10.1016/j.bbabio.2019.06.012.

(31) Griffin, M.; Khan, R.; Basu, S.; Smith, S. Ion Channels as Therapeutic Targets in High Grade Gliomas. Cancers. 2020. 10.3390/cancers12103068.

(32) Jain, A.; Gosling, J.; Liu, S.; Wang, H.; Stone, E. M.; Chakraborty, S.; Jayaraman, P. S.; Smith, S.; Amabilino, D. B.; Fromhold, M.; Long, Y. T.; Pérez-García, L.; Turyanska, L.; Rahman, R.; Rawson, F. J. Wireless Electrical–Molecular Quantum Signalling for Cancer Cell Apoptosis. Nature Nanotechnology 2023 19:1 2023, 19 (1), 106–114. 10.1038/s41565-023-01496-y.

(33) Lee, D.; Lee, D.; Won, Y.; Hong, H.; Kim, Y.; Song, H.; Pyun, J. C.; Cho, Y. S.; Ryu, W.; Moon, J. Insertion of Vertically Aligned Nanowires into Living Cells by Inkjet Printing of Cells. Small 2016, 12 (11). 10.1002/smll.201502510.

(34) Kwon, Y. T.; Kim, Y. S.; Kwon, S.; Mahmood, M.; Lim, H. R.; Park, S. W.; Kang, S. O.; Choi, J. J.; Herbert, R.; Jang, Y. C.; Choa, Y. H.; Yeo, W. H. All-Printed Nanomembrane Wireless Bioelectronics Using a Biocompatible Solderable Graphene for Multimodal Human-Machine Interfaces. Nat Commun 2020, 11 (1). 10.1038/s41467-020-17288-0.

(35) Ryu, H.; Fuwad, A.; Yoon, S.; Jang, H.; Lee, J. C.; Kim, S. M.; Jeon, T. J. Biomimetic Membranes with Transmembrane Proteins: State-of-the-Art in Transmembrane Protein Applications. International Journal of Molecular Sciences. 2019. 10.3390/ijms20061437.

(36) Ren, S.; Zhang, Z.; Dong, Z. Biomimetic Ion Channels: An Emerging and Promising Material for Therapeutic Ion Channelopathies. Trends in Chemistry. Cell Press December 1, 2024. 10.1016/j.trechm.2024.10.005.

(37) Tunuguntla, R. H.; Henley, R. Y.; Yao, Y. C.; Pham, T. A.; Wanunu, M.; Noy, A. Enhanced Water Permeability and Tunable Ion Selectivity in Subnanometer Carbon Nanotube Porins. Science (1979) 2017, 357 (6353). 10.1126/science.aan2438.

(38) Duque, J. G.; Eukel, J. A.; Pasquali, M.; Schmidt, H. K. Self-Assembled Nanoparticle-Nanotube Structures (NanoPaNTs) Based on Antenna Chemistry of Single-Walled Carbon Nanotubes. Journal of Physical Chemistry C 2009, 113 (43). 10.1021/jp906038k.

(39) Warakulwit, C.; Nguyen, T.; Majimel, J.; Delville, M. H.; Lapeyre, V.; Garrigue, P.; Ravaine, V.; Limtrakul, J.; Kuhn, A. Dissymmetric Carbon Nanotubes by Bipolar Electrochemistry. Nano Lett 2008, 8 (2). 10.1021/nl072652s.

(40) Robinson, A. J.; Jain, A.; Rahman, R.; Abayzeed, S.; Hague, R. J. M.; Rawson, F. J. Impedimetric Characterization of Bipolar Nanoelectrodes with Cancer Cells. ACS Omega 2021, 6 (44). 10.1021/acsomega.1c03547.

(41) Ho, N. T.; Siggel, M.; Camacho, K. V.; Bhaskara, R. M.; Hicks, J. M.; Yao, Y. C.; Zhang, Y.; Köfinger, J.; Hummer, G.; Noy, A. Membrane Fusion and Drug Delivery with Carbon Nanotube Porins. Proc Natl Acad Sci U S A 2021, 118 (19). 10.1073/pnas.2016974118.

(42) Sullivan, K.; Zhang, Y.; Lopez, J.; Lowe, M.; Noy, A. Carbon Nanotube Porin Diffusion in Mixed Composition Supported Lipid Bilayers. Sci Rep 2020, 10 (1). 10.1038/s41598-020-68059-2.

(43) Hicks, J. M.; Yao, Y. C.; Barber, S.; Neate, N.; Watts, J. A.; Noy, A.; Rawson, F. J. Electric Field Induced Biomimetic Transmembrane Electron Transport Using Carbon Nanotube Porins. Small 2021, 17 (32). 10.1002/smll.202102517.

(44) Ji, S. rong; Liu, C.; Zhang, B.; Yang, F.; Xu, J.; Long, J.; Jin, C.; Fu, D. liang; Ni, Q. xing; Yu, X. jun. Carbon Nanotubes in Cancer Diagnosis and Therapy. Biochimica et Biophysica Acta - Reviews on Cancer. 2010. 10.1016/j.bbcan.2010.02.004.

(45) Pavlov, V. A.; Tracey, K. J. Bioelectronic Medicine: Updates, Challenges and Paths Forward. Bioelectronic Medicine. 2019. 10.1186/s42234-019-0018-y.

(46) Tunuguntla, R. H.; Escalada, A.; A Frolov, V.; Noy, A. Synthesis, Lipid Membrane Incorporation, and Ion Permeability Testing of Carbon Nanotube Porins. Nat Protoc 2016, 11 (10). 10.1038/nprot.2016.119.

(47) Moreira, L.; Fulchiron, R.; Seytre, G.; Dubois, P.; Cassagnau, P. Aggregation of Carbon Nanotubes in Semidilute Suspension. Macromolecules 2010, 43 (3), 1467–1472. 10.1021/MA902433V/ASSET/IMAGES/MEDIUM/MA-2009-02433V_0003.GIF.

(48) Dubey, R.; Dutta, D.; Sarkar, A.; Chattopadhyay, P. Functionalized Carbon Nanotubes: Synthesis, Properties and Applications in Water Purification, Drug Delivery, and Material and Biomedical Sciences. Nanoscale Adv 2021, 3 (20), 5722–5744. 10.1039/D1NA00293G.

(49) Phospholipid spherules (liposomes) as a model for biological membranes - PubMed. https://pubmed.ncbi.nlm.nih.gov/5646182/ (accessed 2025-04-16).

(50) Djamgoz, M. B. A.; Mycielska, M.; Madeja, Z.; Fraser, S. P.; Korohoda, W. Directional Movement of Rat Prostate Cancer Cells in Direct-Current Electric Fieldinvolvement of Voltagegated Na+ Channel Activity. J Cell Sci 2001, 114 (14), 2697–2705. 10.1242/JCS.114.14.2697.

(51) Melikov, R.; De Angelis, F.; Moreddu, R. High-Frequency Extracellular Spiking in Electrically-Active Cancer Cells. March 17, 2024. 10.1101/2024.03.16.585162.

(52) Habib-E-Rasul Mullah, S.; Komuro, R.; Yan, P.; Hayashi, S.; Inaji, M.; Momose-Sato, Y.; Loew, L. M.; Sato, K. Evaluation of Voltage-Sensitive Fluorescence Dyes for Monitoring Neuronal Activity in the Embryonic Central Nervous System. Journal of Membrane Biology 2013, 246(9). 10.1007/s00232-013-9584-1.

(53) Olszewski, D. Asymmetry Index for Data and Its Verification in Dimensionality Reduction and Data Visualization. Inf Sci (N Y) 2025, 689, 121405. 10.1016/J.INS.2024.121405.

(54) Moini, J.; Avgeropoulos, N. G.; Samsam, M. Cytology of the Nervous System. Epidemiology of Brain and Spinal Tumors 2021, 41–63. 10.1016/B978-0-12-821736-8.00012-1.

(55) Moini, J.; LoGalbo, A.; Ahangari, R. Neural Tissue. Foundations of the Mind, Brain, and Behavioral Relationships 2024, 39–62. 10.1016/B978-0-323-95975-9.00017-2.

(56) Blackiston, D. J.; McLaughlin, K. A.; Levin, M. Bioelectric Controls of Cell Proliferation: Ion Channels, Membrane Voltage and the Cell Cycle. Cell Cycle. 2009. 10.4161/cc.8.21.9888.

(57) Fialova, J. L. Novel Mitochondria-Targeted Drugs for Cancer Therapy. Mini-Reviews in Medicinal Chemistry 2022, 21 (7). 10.2174/18755607mtexxnjqhy.

(58) Spannl, S.; Buhl, T.; Nellas, I.; Zeidan, S. A.; Iyer, K. V.; Khaliullina, H.; Schultz, C.; Nadler, A.; Dye, N. A.; Eaton, S. Glycolysis Regulates Hedgehog Signalling via the Plasma Membrane Potential. EMBO J 2020, 39 (21). 10.15252/embj.2019101767.

(59) Vasey, C. E.; Cavanagh, R. J.; Taresco, V.; Moloney, C.; Smith, S.; Rahman, R.; Alexander, C. Polymer Pro-Drug Nanoparticles for Sustained Release of Cytotoxic Drugs Evaluated in Patient-Derived Glioblastoma Cell Lines and In Situ Gelling Formulations. Pharmaceutics 2021, Vol. 13, Page 208 2021, 13 (2), 208. 10.3390/PHARMACEUTICS13020208.

(60) Molenaar, R. J. Ion Channels in Glioblastoma. ISRN Neurol 2011, 2011. 10.5402/2011/590249.

(61) Gensbittel, V.; Kräter, M.; Harlepp, S.; Busnelli, I.; Guck, J.; Goetz, J. G. Mechanical Adaptability of Tumor Cells in Metastasis. Developmental Cell. 2021. 10.1016/j.devcel.2020.10.011.

