## Supporting information for "Bioelectronic Modulation of Glioblastoma via Wireless Carbon Nanotube Porin Interfaces"

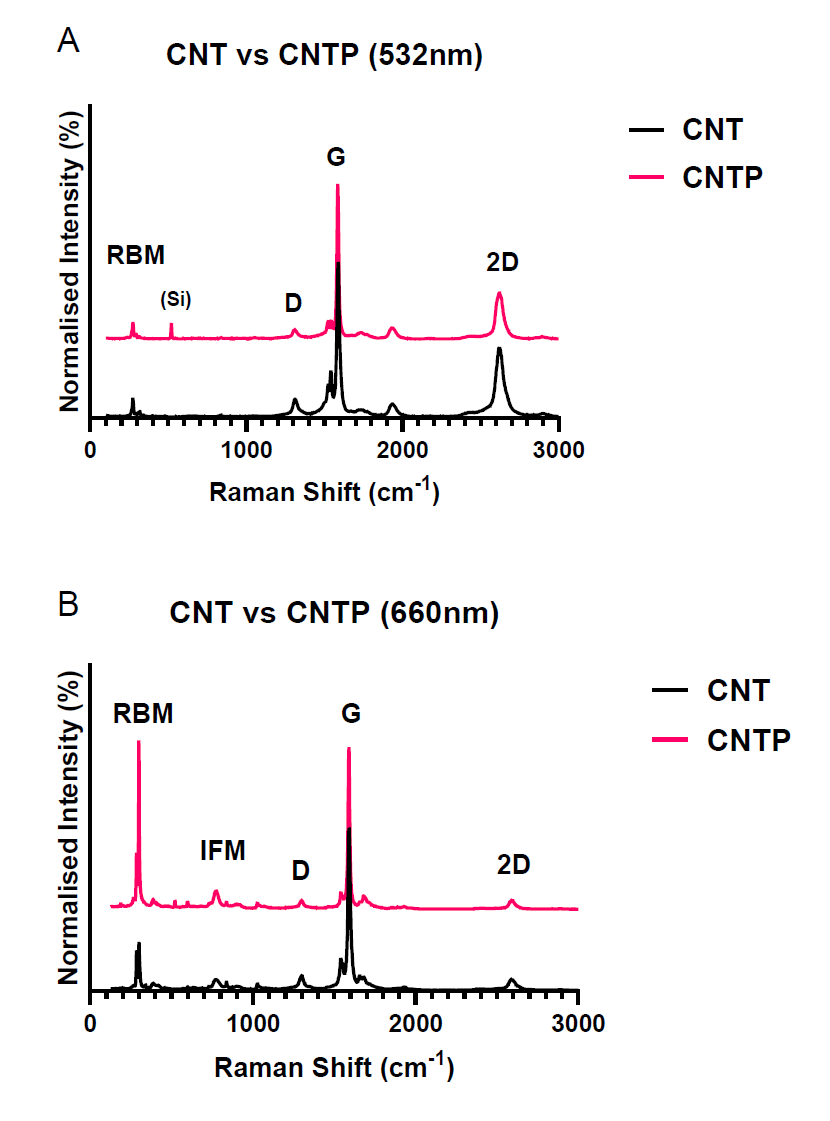

**Figure S1.** A) 532 nm and B) 660 nm Raman spectra of CNT and CNTP samples.

The Raman spectra of CNTs shown in Figures S1A&B are consistent with expectation and evidence the characteristic RBM, D, G and 2D bands known for single-walled carbon nanotubes. It is important to note that the spectra collected do not describe all nanotubes present within the sample, rather the subset of nanotubes enhanced through resonance effects (here these are a set of specific chirlaities of semiconducting nanotubes visible in the spectra due to the correct match of the excitation laser to the E_22_^S^ optical transition). There is no direct evidence for the porin in the spectra of CNTPs, i.e., we do not see Raman bands of the porin, due to the previously mentioned resonant enhancement of nanotube bands. However, the reduction in the intensity of CNT D and 2D bands (relative to the G band) in the 532 and 660 nm Raman spectra of CNTPs provides tentative evidence for the presence of the porin, with lipid-nanotube interactions after surface passivation likely mediating debundling of CNT aggregates. An increase in the intensity of one of the RBMs (relative to the G band) in the 660 nm spectrum of CNTPs supports this assertion.

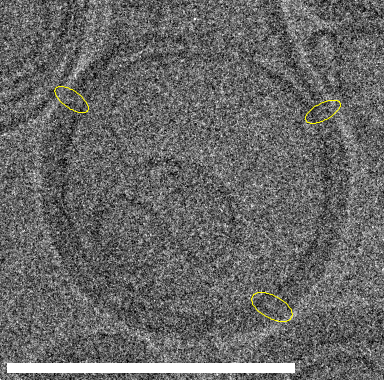

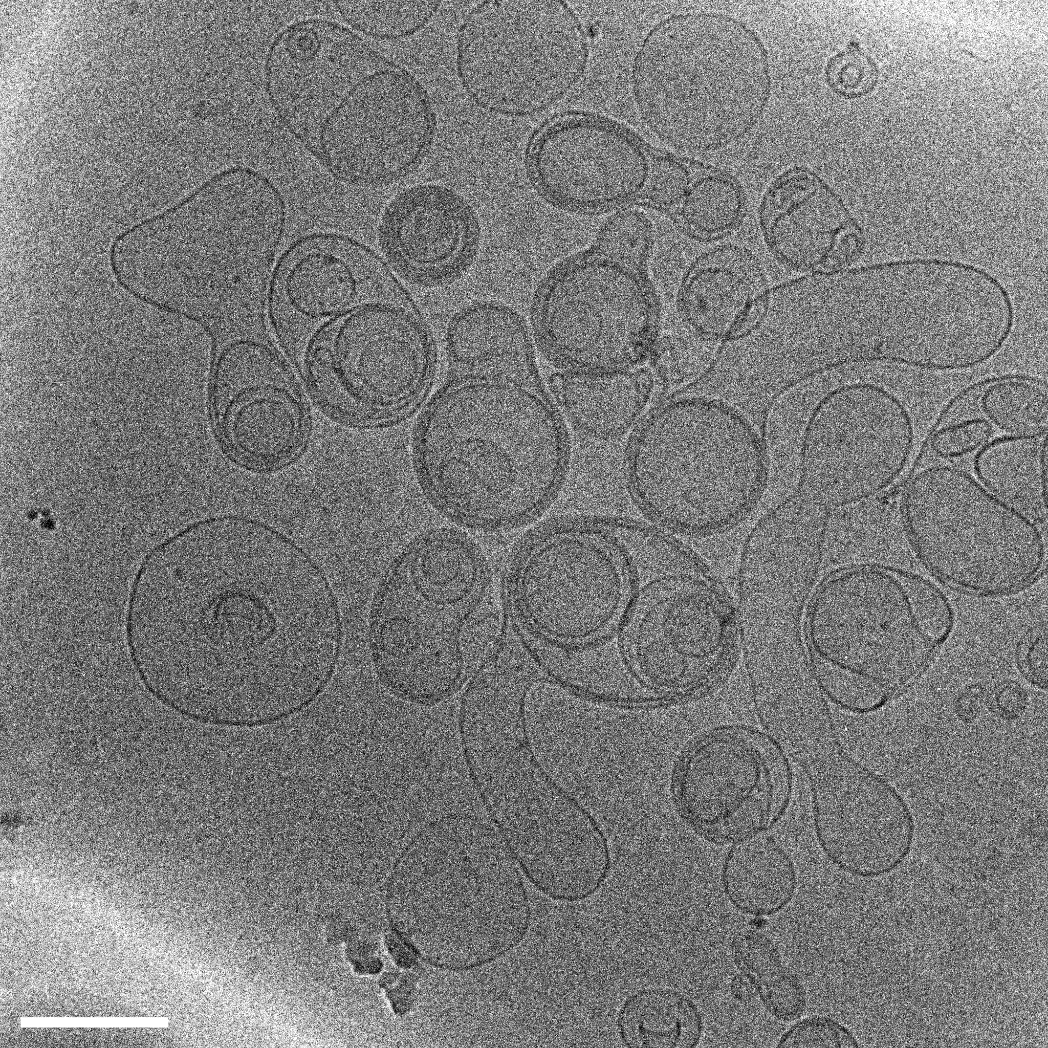

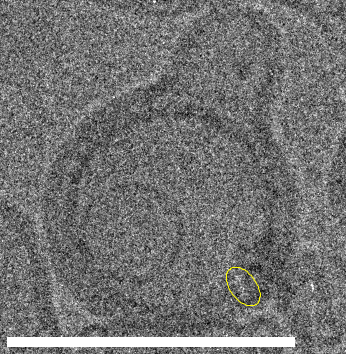

**A**

**B**

**C**

**Figure S2.** cryo-TEM images of GUVs with CNTPs within the membrane. Yellow boxes in A) correspond to zoomed in areas shown in B) (lower) and C) (upper), respectively. Scale bars are 200 nm. CNTPs visible in B) and C) within the membrane are circled.

CNTP structures in S2B, top left found to be 22.0nm x 2.8 nm, top right measured at 24.5 nm x 3.1 nm and bottom right recorded to be 24.5 nm x 2.4 nm. The CNTP structure circled in S2C are measured at 26.1 nm x 2.5 nm. Due to intrinsic technical limitations of TEM to measure size and the variability in the lengths of the CNTs within a sample, previous reports (Tunuguntla et al., 2016) indicate that CNTPs likely measure ~10 x 0.7 nm, so the measurements observed here broadly support the incorporation of porins into the membranes of GUVs.

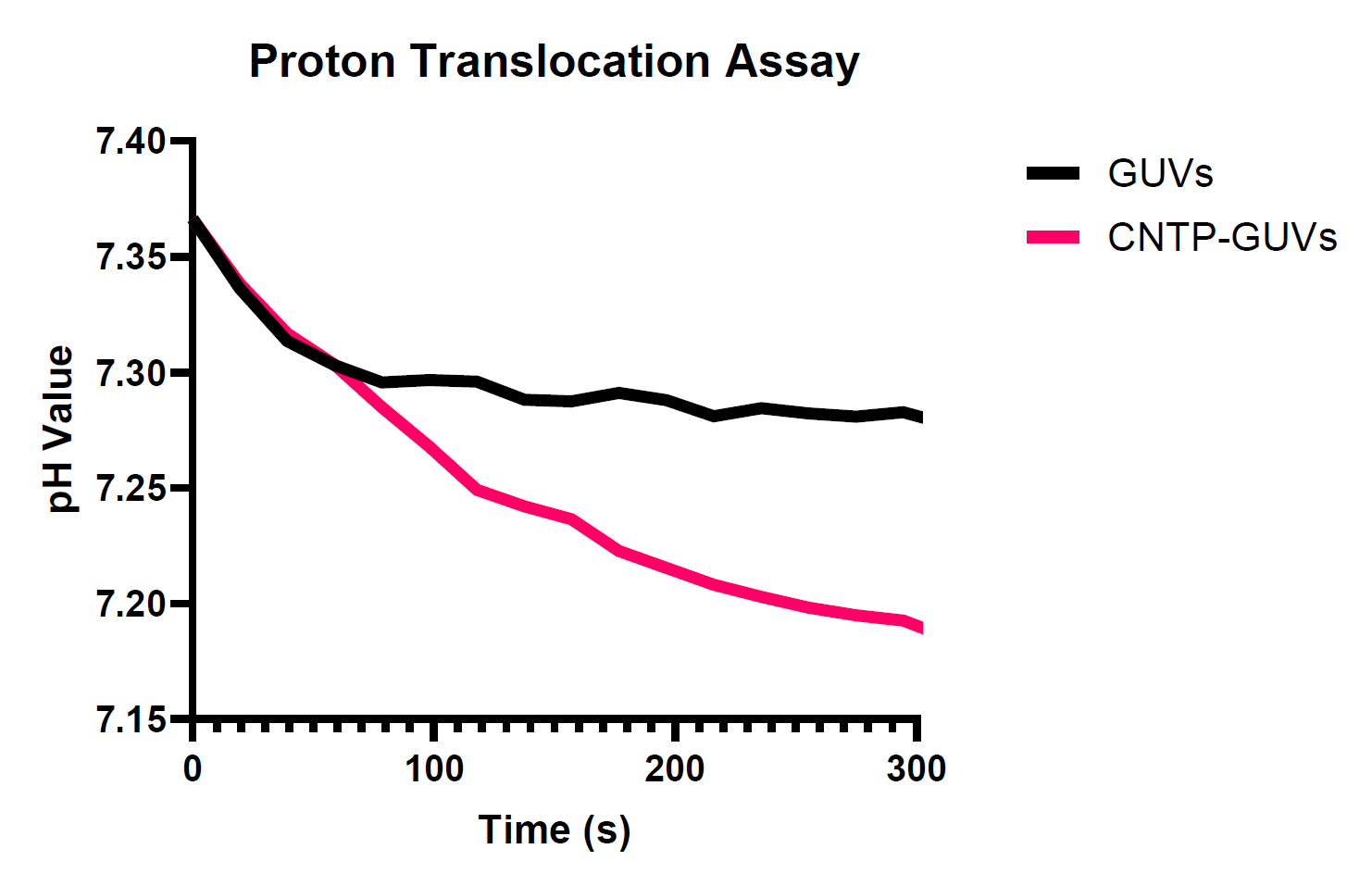
**Figure S3.** Proton translocation assay of GUVs and CNTP-GUVs read at excitation/emission wavelengths of 448/516 nm, converted to pH values, over time.

The notable decrease in pH observed in CNTP-GUVs compared to GUVs demonstrates that lipid bilayers are permeable to protons and can gradually equilibrate a pH gradient across the membrane. The CNTPs can serve as a useful tool for monitoring that gradient.

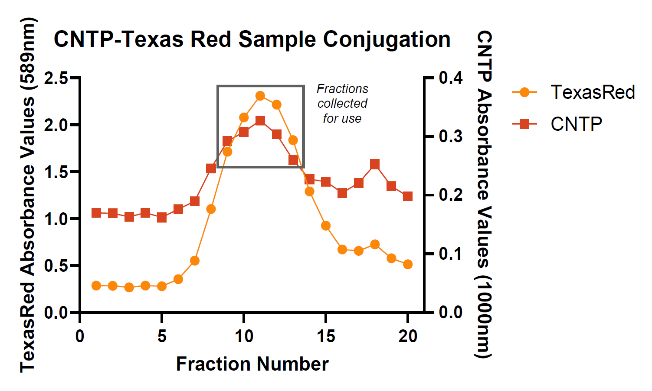

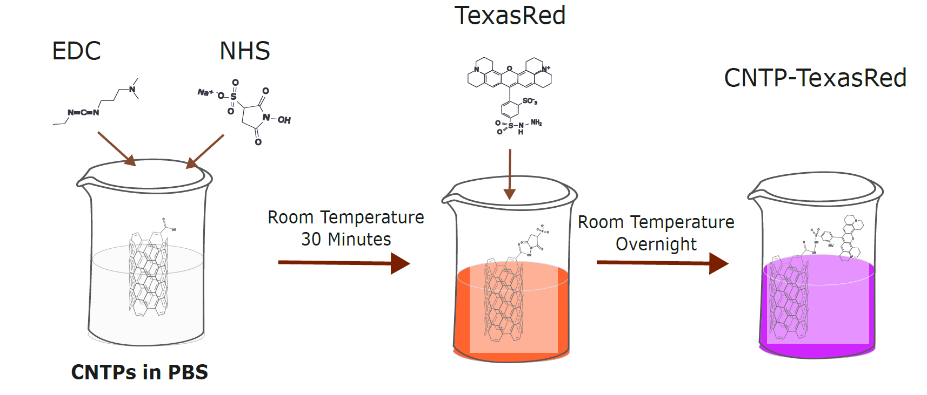

**A**

**B**

**Figure S4.** A) CNTP modification with TexasRed via EDC/NHS filtered through Sepharose column with fractions collected (9-13) and absorbances measured at 589 nm for Texas-Red and 1000 nm for CNTPs. B) Schematic of the reaction of CNTPs with TexasRed via EDC/NHS chemistry. Solubilization of CNTPS with EDC (80 mm) and NHS (20 mm) which is left to react at room temperature for 30 minutes. Fluorophores TexasRed are then added and left overnight to react in the dark at room temperature.

Texas Red (16mM) was conjugated with CNTPs following an EDC/NHS coupling chemistry protocol simiarly to previous work (Hicks et al., 2021) shown in figure S4B. The obtained material was filtered through a fresh Sepharose CL-6B column, and the fractions werer collected. Those with an absorbances at both 589 nm (Texas Red) and 1000 nm (CNTPs) suggesting successful conjugation.

**Figure S5.** Confocal microscopy 3D orthogonal z-stack confirming CNTPs within the plasma membrane in GCE-31 (left) and GIN-31 (right). Scale bar =100µm

XZ

YZ

YZ

XZ

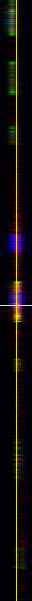

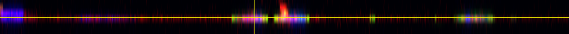

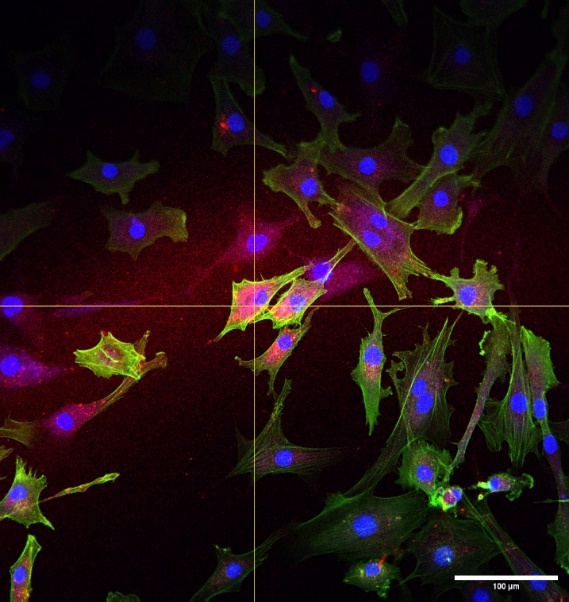

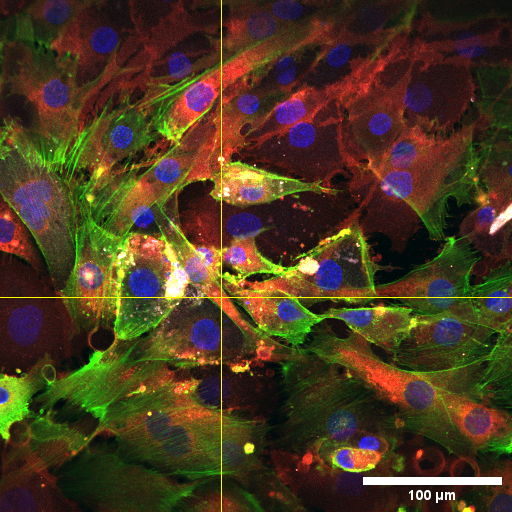

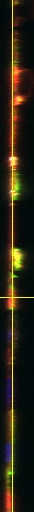

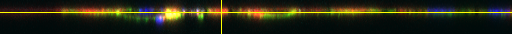

GCE-31

GIN-31

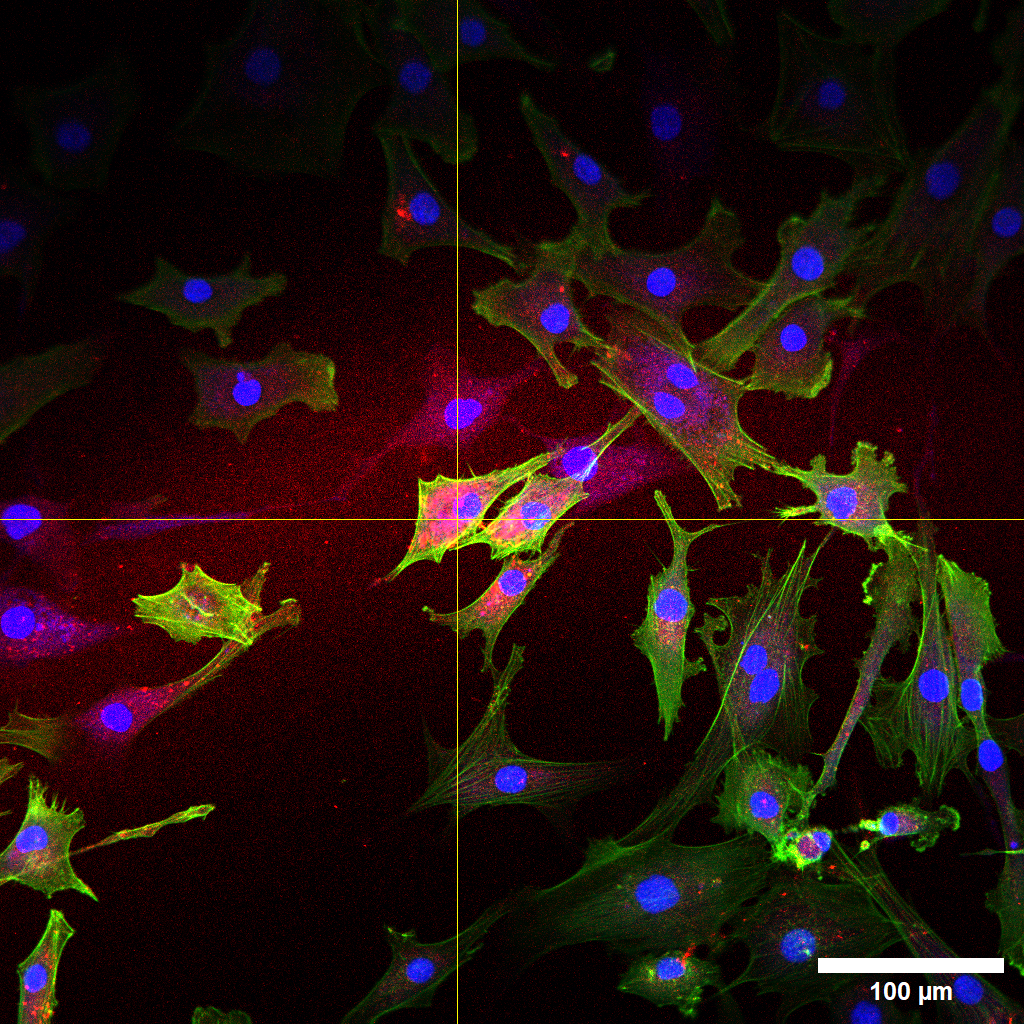

Cells were incubated with CNTP-TexasRed for 4 h and then fixed with paraformaldehyde followed by staining with Actin Phalloidin (Alexa 488 green) and Hoechst nuclear (blue). CNTP-TexasRed (red) is seen in the cytoplasm. The bright red colouration indicates the presence of CNTPS, and the Z-stacks confirm its location within the cell membrane.

**Figure S6**. Raw Data Time-dependent changes in V_mem_ (cf. fluorescence intensity) within GBM cells, GCE-31 and GIN-31, with and without CNTP-TexasRed across different applied voltages from 0 to 2.5V. A) and B) GCE-31 cells without and with CNTPs, respectively. C) and D) GIN-31 cells without and with CNTPs, respectively. Normalised cell fluorescence was extracted via ImageJ and used to quantify the percentage change (*ΔF/F)* over time. N=2, n=10.

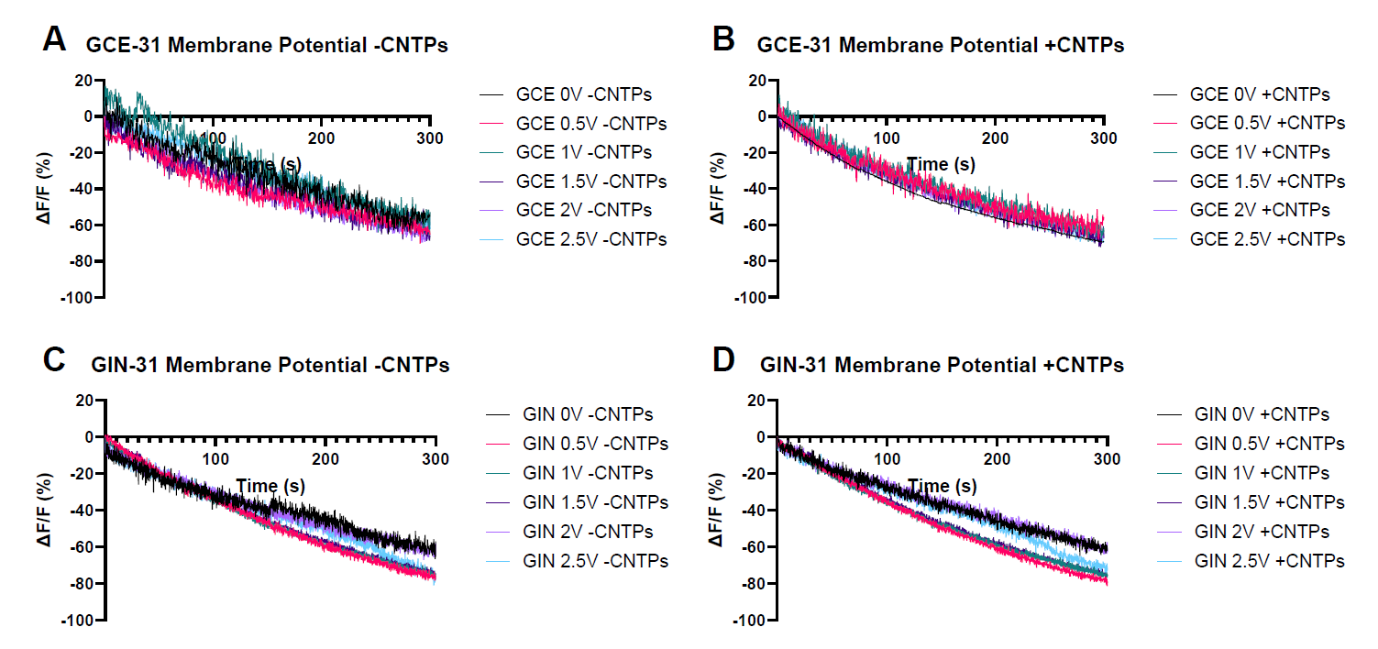

Figure S7 presents a visual representation of the rate of change in the V_mem_ signal across different applied voltages, both with and without CNTPs.This cn be further confirmed through the calculated metrics of the rate of change, such as the peak rate of change, trough rate of change and asymmetry index, which can be found in Table S2.

**Figure S7** Raw Rate of change plots for GCE-31 and GIN-31 have been generated using the dataset from Figure S6. Python was applied with a Gaussian smoothing filter to make the trends clearer. A) and B) GCE-31 cells without and with CNTPs, respecitvely. C) and D) GIN-31 cells without and with CNTPs, respectively.

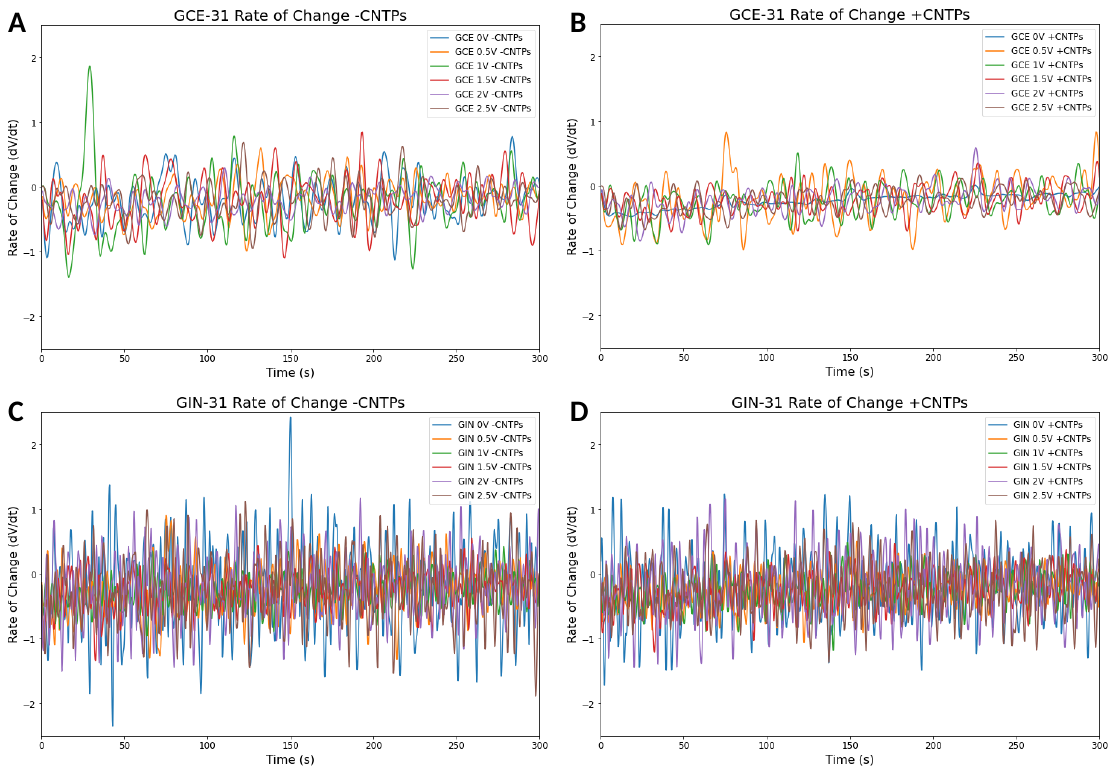

**Table S1**. Metrics of V_mem_ signal characteristics in GCEs and GINs such as the Peak Rate of Change – Speed of changes of peaks, Trough Rate of Change - Speed of changes of troughs, Asymmetry Index – Peak/troughs and Spike Rate (Hz) – Number of spikes per second

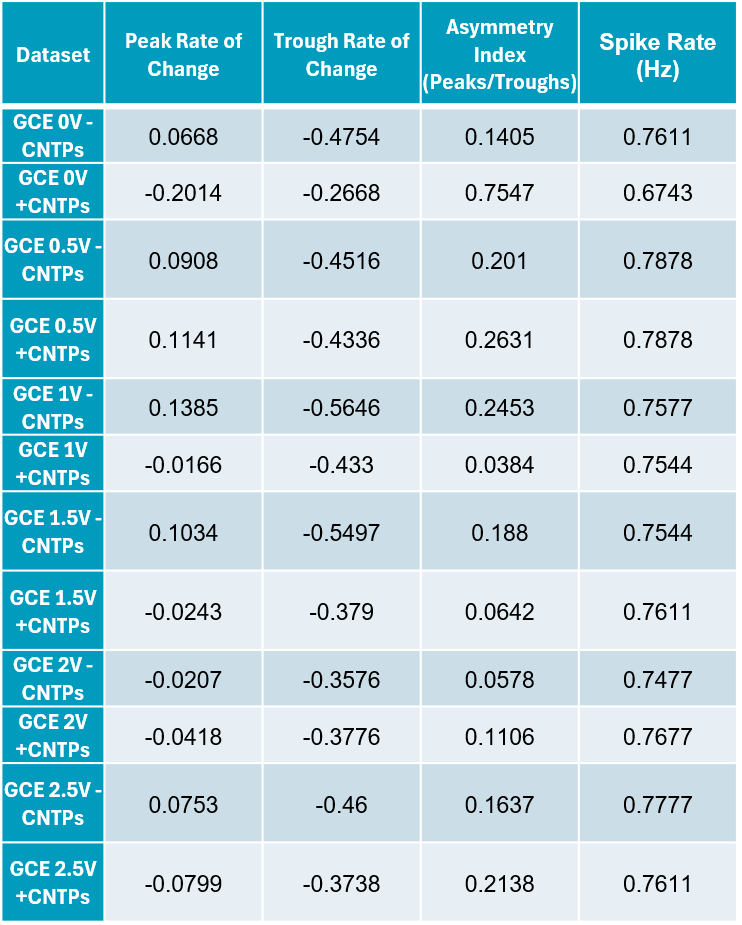

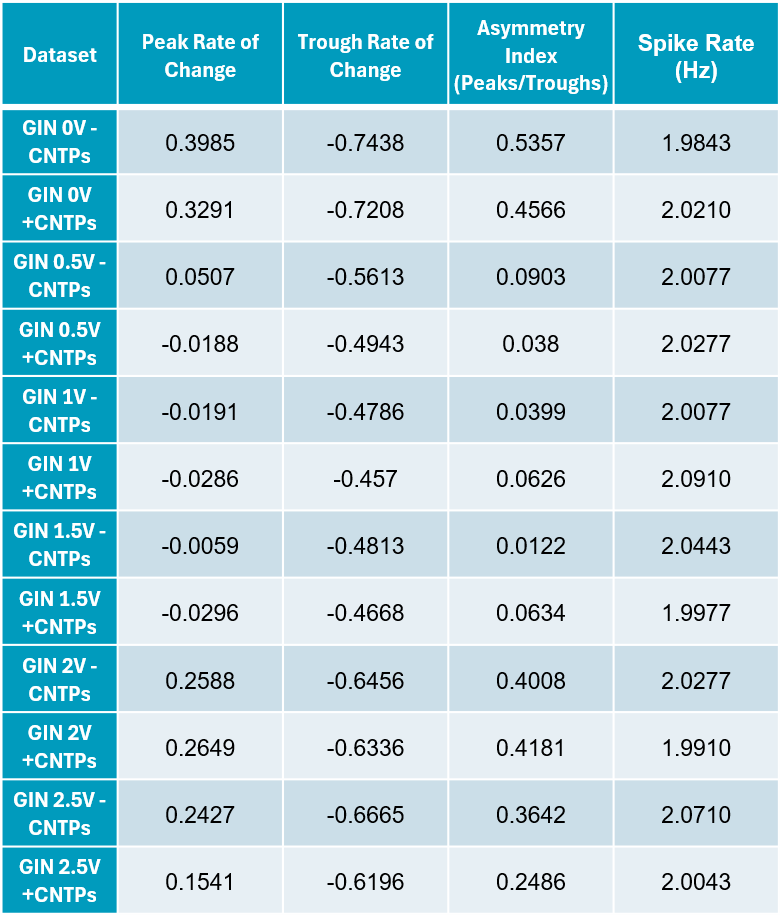

**Figure S8**. Flow cytometry cell cycle gating strategy.

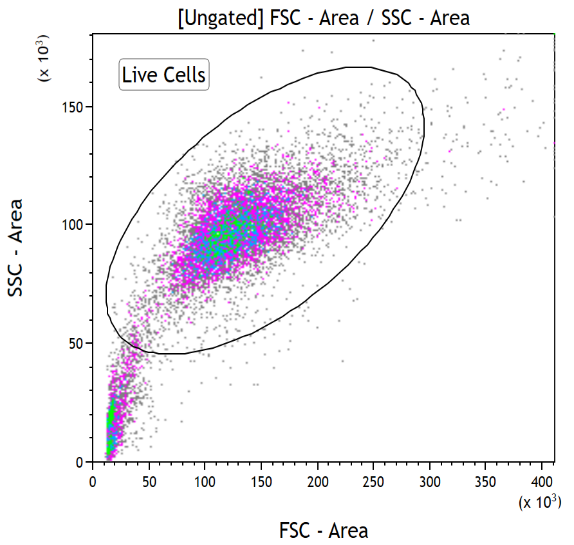

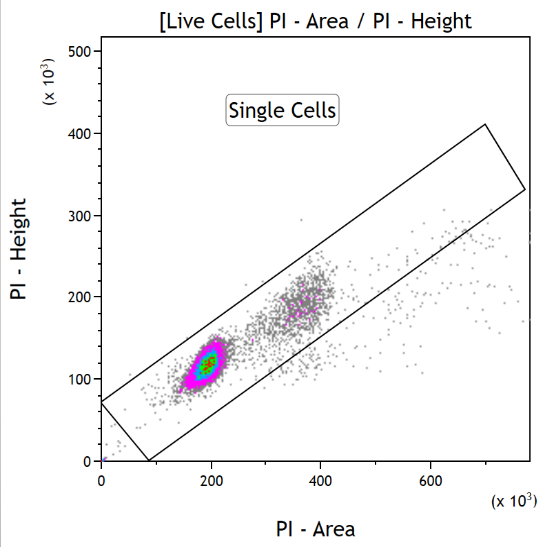

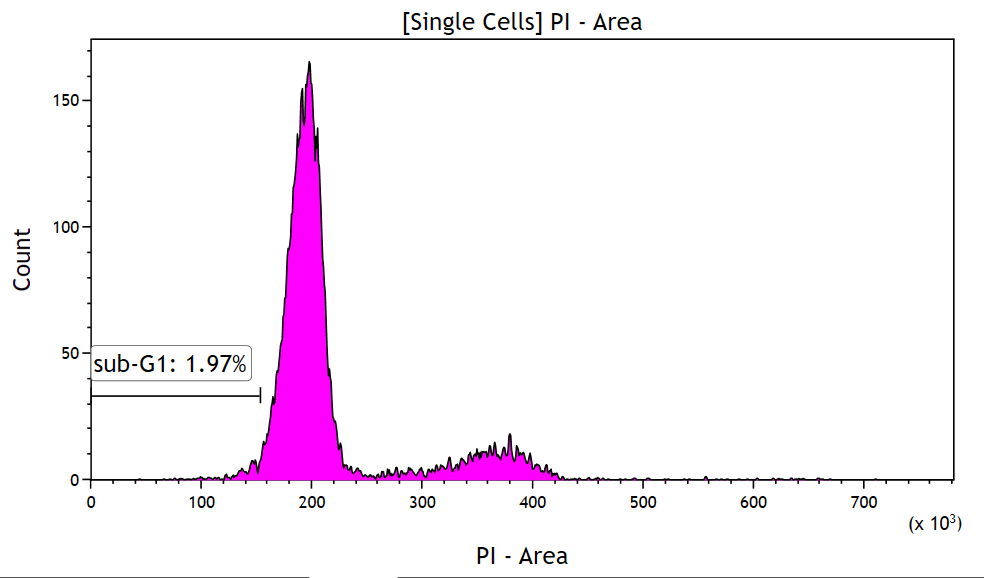

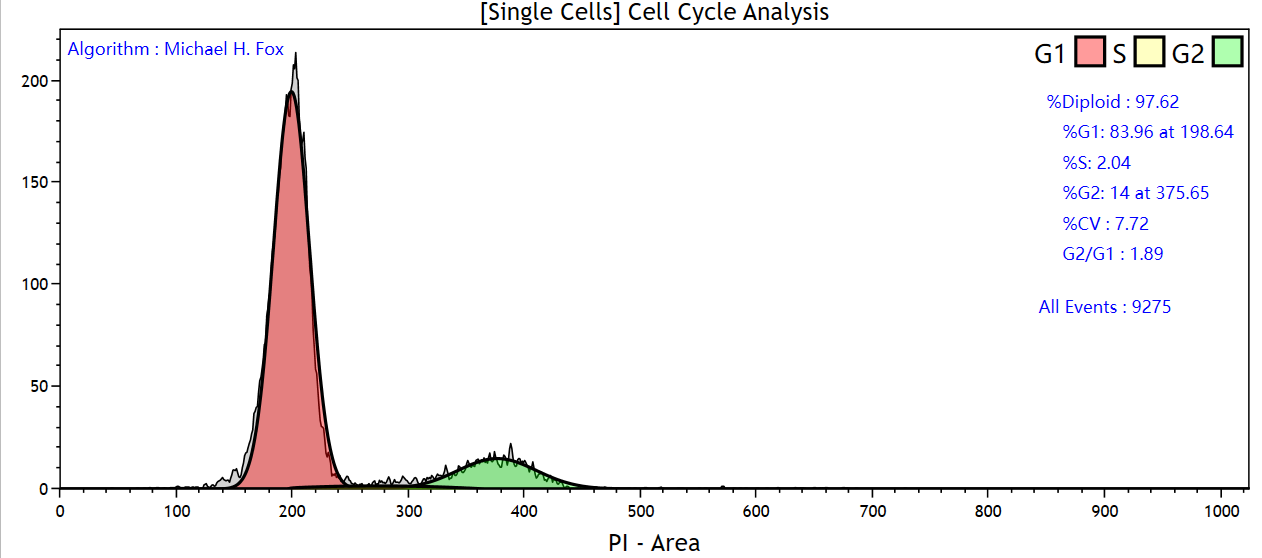

Cells post-exposure to conditions after their respective time point was fixed with ethanol which was then treated with FxCycle™ PI/RNAse staining solution prior to data acquisition via flow cytometry. The data was analysed with Kaluza Analysis 2.1. Live cells were selected using Forward Scattering (FSC) vs Side Scattering (SSC) gating to avoid doublets. Single cells were gated using PI - Area vs PI - Height and analyzed via the software's cell cycle algorithm to display cell cycle populations. If %Diploid was below 100, manual gating was employed to identify the sub-G1 phase.

METHODS

**Carbon Nanotube Porin Synthesis** – Protocol was adapted from a previous publication from the Noy group^35^. 28mg of (1,2-dioleoylsn-glycero-3-phosphocholine) DOPC (Avanti Polar Lipids) were dried within a scintillation vial in a Vacuum Centrifuge Evaporator (Labconco). Meanwhile 0.5mg of single-walled CNTS (SWCNTs) (773735, Sigma Aldrich) were baked incrementally from 100 ºC, which is increased by 20 ºC every 2 minutes until a final temperature of 475ºC. The purified CNTs were added to 14ml of Milli-Q purified H_2_O and lipids from dried vial and placed in a sonicator water bath (Fisherbrand) for approximately 30 minutes. Once solubilized, the 14ml solution was then pipetted into another 20ml glass scintillation vial and transferred to a microbiological safety cabinet. The cabinet included the ultrasonicator (Qsonica) within a sound enclosure with a microtip diameter of 0.25 inches. The sample was capped by a custom-made hole of 0.45 inches in diameter and placed in a custom sample holder that permitted water circulation within the holder at a constant temperature of 25°C from a recirculation chiller (Polyscience). The ultrasonicator was then programmed to sonicate the sample for 16 hours in pulses of 3 seconds on and 1 second off, at an amplitude of 35%. When the sonication had been completed, 1mL of the sample was then pipetted into 1.5ml microcentrifuge tubes, which were then placed in a micro centrifuge for 1 hour at 10000 rcf. The supernatant was then carefully pipetted and placed into a 15ml falcon tube.

**CNTP Modification –** Protocol was obtained and adapted from previous publication within the Rawson group^41^_._ 1 ml of pre-made CNTPs was pipetted into a microcentrifuge tube and spun in a microcentrifuge at 14000 x G rpm for 6 minutes. The supernatant was then aspirated and placed into an 8 ml scintillation vial and dried in a Rotavapor Evaporator until a film is formed. During this period, solutions of 6-aminofluorescein (6AF) (Merck, 201634) or TexasRed Hydrazide (Stratech, 481-AAT). The 6AF solution was made up from 25mM 6-aminofluorescein or 10mM of TexasRed dissolved in a 50:50 ratio of 99% Ethanol and Phosphate-Buffered Saline Buffer (PBS) of 7.4 pH. The CNTPS were then re-suspended in a 80mM 1-ethyl-3-(-3-dimethylaminopropyl) carbodiimide hydrochloride (EDC) / 20mM N-hydroxysuccinimide (NHS) carbodiimide (NHS) solution and sonicated briefly in a water bath for 1 minute, the solution would be cloudy and slightly dark in colour. The solution was then left on the benchtop at room temperature for 15 minutes. The Texas Red or 6AF solution was then spun on the microcentrifuge using the “short” function at max speed of 16000 x g for 30 seconds. Once finished 100ul was pipetted into the CNTP EDC/NHS solution, wrapped in foil and then placed in a drawer in the dark and left overnight at room temperature. The following day, the solution would have turned dark orange. A sepharose column onto a stand and allowing it to warm to room temperature. Meanwhile, water was drained in the column by uncapping and flushing the column with 5ml with PBS. A 96 well plate (Nunc) was placed underneath, with the well A1 directly underneath the tip at the bottom of the column. The CNTP-6AF/CNTP-Red solution was then pipetted into the column and once the solution could be visibly seen 1cm within the gel bed of the column, fractions were collected. 10 drops were collected per well, refilling the top of the column with PBS when necessary and continued to collect fractions until the lighter orange colour has been collected. The fractions were then analysed with a TECAN Plate reader, with the Absorbance at 1000nm for CNTPs, 495nm for 6AF and 589nm for Texas Red.

**Raman Spectroscopy**

Micro Raman spectroscopy was performed using a HORIBA LabRAM HR Raman microscope. Spectra were acquired using both a 532 and 660 nm laser (at <0.5 mW power), a 100x objective and a 200 μm confocal pinhole. To simultaneously scan a range of Raman shifts, 600 lines mm^-1^ rotatable diffraction gratings along a path length of 800 mm were employed. Spectra were detected using a Synapse CCD detector (1024 pixels) thermoelectrically cooled to −60 °C. Before spectra collection, the instrument was calibrated using the zero-order line and a standard Si(100) reference band at 520.7 cm^-1^. The spectral resolution is better than 1.8 and 1.2 cm^-1^ in these configurations, respectively. Samples were drop cast onto silicon wafers, and dried under ambient conditions, for analysis.

**Cryo-TEM**

Samples were prepared using a Gatan CP3 Cryoplunge providing a controlled environment (70 – 80 % humidity, 18 oC), by depositing 3 μL of sample onto a TEM grid (300 mesh Cu, holey carbon or holey carbon/graphene oxide support film, EM Resolutions Ltd). Samples were blotted (1.5 s) before plunging into liquid ethane to vitrify. Samples were maintained under liquid nitrogen (-196 oC) until transfer to a TEM cryo sample holder (Gatan 926) and held at or below -160ºC during analysis (Gatan Smartset 900). TEM images were recorded on JEOL 2100 Plus, operating at 200 kV using a Gatan Ultrascan 100XP camera. CNTP sizes were acquired via ImageJ.

**FTIR**

CNTP-TexasRed was centrifuged in microcentrifuge tube for 1 hour, supernatant was then removed and freeze-dried at -80ºC for over 24 hourswhich was then analysed through an FTIR Spectrometer (Cary 530, Agilent) at transmittance from 4000 – 650 cm^-1^.

**Confocal Microscopy**

Cells were seeded at 1x10^5^ cells/well in a 24 well plate (Costar), left to incubate overnight. The following day, the media was aspirated from the well to be replaced 20µL of CNTP-Texas Red or CNTPs was added to 80µL of DMEM incubated for at minimum 4 hours. Cells were then stained with an Actin (23115, AAT Bioquest, Stratech) and NucBlue according to their recommended protocol. Cells were then fixed with 4% paraformaldehyde for 30 mins, before being washed with PBS prior to imaging with a Leica TCS SPE confocal microscope.

**Proton Translocation Assay**

Three buffers (10 mM HEPES-K, 150 mM NaCl, 30 mM KCl, 500 ml dH2O) were prepared and labeled A, B, and C at pH ~9.4. Buffer A received 10 mM pyranine; pH for A and B was adjusted to 7.51, while C was adjusted to 6.9 using 1M HCl. DOPC (80 µl) was dried in vials; 1 ml of CNTPs was centrifuged, then supernatant dried and rehydrated in Buffer A. Supernatants from rehydrated CNTPs were combined with the DOPC vials and sonicated. After 30 min at room temp for vesicle formation, the microcentrifuge tubes were immersed in liquid nitrogen (LN2) for 45 seconds for 10 cycles. Solutions were extruded through an Avanti Mini Extruder. Buffer B was added to remove impurities before the extruded solution was collected in a 96-well plate from a size exclusion column. Clear fractions were analyzed via fluorescence (448 nm/516 nm) with Buffer C and HCl added sequentially
